# Neuronal Synchrony and Critical Bistability: Mechanistic Biomarkers for Localizing the Epileptogenic Network

**DOI:** 10.1101/2023.05.21.541570

**Authors:** Sheng H Wang, Gabriele Arnulfo, Lino Nobili, Vladislav Myrov, Paul Ferrari, Philippe Ciuciu, Satu Palva, J Matias Palva

## Abstract

**Objective:** Post-surgical seizure freedom in drug-resistant epilepsy (DRE) patients varies from 30 to 80%, implying that in many cases the current approaches fail to fully map the epileptogenic zone (EZ). This suggests that the EZ entails a broader epileptogenic brain network (EpiNet) beyond the seizure-zone (SZ) that show seizure activity.

**Methods:** We first used computational modeling to identify putative complex-systems- and systems-neuroscience-driven mechanistic biomarkers for epileptogenicity. We then extracted these epileptogenicity biomarkers from stereo-EEG (SEEG) resting-state data from DRE patients and trained supervised classifiers to localize the SZ with these biomarkers against gold-standard clinical localization. To further explore the prevalence of these pathological biomarkers in an extended network outside of the clinically-identified SZ, we also used unsupervised classification.

**Results:** Supervised SZ-classification trained on individual features achieved accuracies of 0.6–0.7 areaunder-the-receiver-operating-characteristics curve (AUC). However, combining all criticality and synchrony features improved the AUC up to 0.85.

Unsupervised classification uncovered an EpiNet-like cluster of brain regions with 51% of regions outside of SZ. Brain regions in this cluster engaged in inter-areal hypersynchrony and locally exhibited high amplitude bistability and excessive inhibition, which was strikingly similar to the high seizure-risk regime revealed by computational modeling.

**Significance:** The finding that combining biomarkers improves EZ localization shows that the different mechanistic biomarkers of epileptogenicity assessed here yield synergistic information. On the other hand, the discovery of SZ-like pathophysiological brain dynamics outside of the clinically-defined EZ provides experimental localization of an extended EpiNet.

**Key points:**

- We advanced novel complex-systems- and systems-neuroscience-driven biomarkers for epileptogenicity
- Increased bistability, inhibition, and power-low scaling exponents characterized our model operating in a high seizure-risk regime and SEEG oscillations in the seizure-zone (SZ)
- Combining all biomarkers yielded more accurate supervised SZ-classification than using any individual biomarker alone
- Unsupervised classification revealed more extended pathological brain networks including the SZ and many non-seizure-zone areas that were previously considered healthy

## 1 Introduction

Epilepsy is an umbrella term for a number of syndromes sharing a common characteristic: an enduring predisposition to having seizures [1, 2]. An epileptic seizure refers to an array of abnormal neuronal activities and hypersynchrony [3, 4] often accompanied with involuntary changes in behaviors, subjective experiences, or loss of consciousness in severe cases [1, 2]. 30–40 % of epilepsy patients have drug-resistant epilepsy (DRE) where anti-seizure medications fail to stop the seizures. The last resort treatment for seizure control in DRE is surgery [5] that aims to resect or disconnect the brain tissue responsible for seizure generation, commonly known as the epileptogenic zone (EZ). Nonetheless, uncertain post-surgical attainment of seizure-freedom (30 – 80%) [6, 7] implies that the biomarkers in current clinical use fail to fully localize the EZ in many patients [8, 9, 10]. New approaches to deepen the understanding of the systems-level mechanisms of epileptogenicity and their individual expression are thus urgently needed.

The EZ-localization is typically assisted by a range of multi-modal mechanistic characteristics, such as MRI-visible lesions [11], metabolic anomalies [12], and neurophysiological anomalies including high gamma oscillations [13], spikes [10], or concurring pathological slow and fast activity [14]. In DRE patients with focal seizure onset, seizure activity is often limited to a small number of brain areas [15]. The identification of the hypothetical EZ involves the visual inspection of SEEG traces during seizure onset and propagation, while considering the behavioral symptoms[15]. However, in light of evidence suggesting that the existence of a broader connected epileptiform network portends poor surgical outcomes, it has been increasingly recognized that epilepsy is a brain-network disease rather than attributable to isolated lesions [16, 17, 18, 19, 20].

Inter-areal synchrony in large-scale brain networks is thought to be instrumental for neuronal communication [21, 22]. Previous SEEG studies have showed that during inter-ictal periods, pathological regions are more synchronized [23]. We have recently reported synchrony levels in the human brain to be predicted by individual positions in a critical state-space [24] so that synchrony among brain regions is moderate in healthy adults but more elevated in epilepsy patients. These evidence supports the hypothesis of criticality being a hallmark of healthy brain functioning and a shift towards super-criticality being characteristic to epileptogenicity.

The *brain criticality* hypothesis posits that healthy brains operate near a phase transition between asynchrony and synchrony [25, 26, 24]. Hallmarks of brain criticality include moderate levels of synchrony (Fig 1, emergent scale-invariant long-range temporal correlations (LRTCs) [27], balanced excitation and inhibition [28, 29], and a unimodal synchrony distribution (Bottom, Fig 1c). Deviations from criticality – either to inadequate (subcriticality) or to excessive synchrony (super-criticality) – have been associated with brain disorders[30]. In particular, neuronal oscillations in epileptic brains has been found to show signs of super-critical-like dynamics [31, 32, 24] .

**Figure 1.**
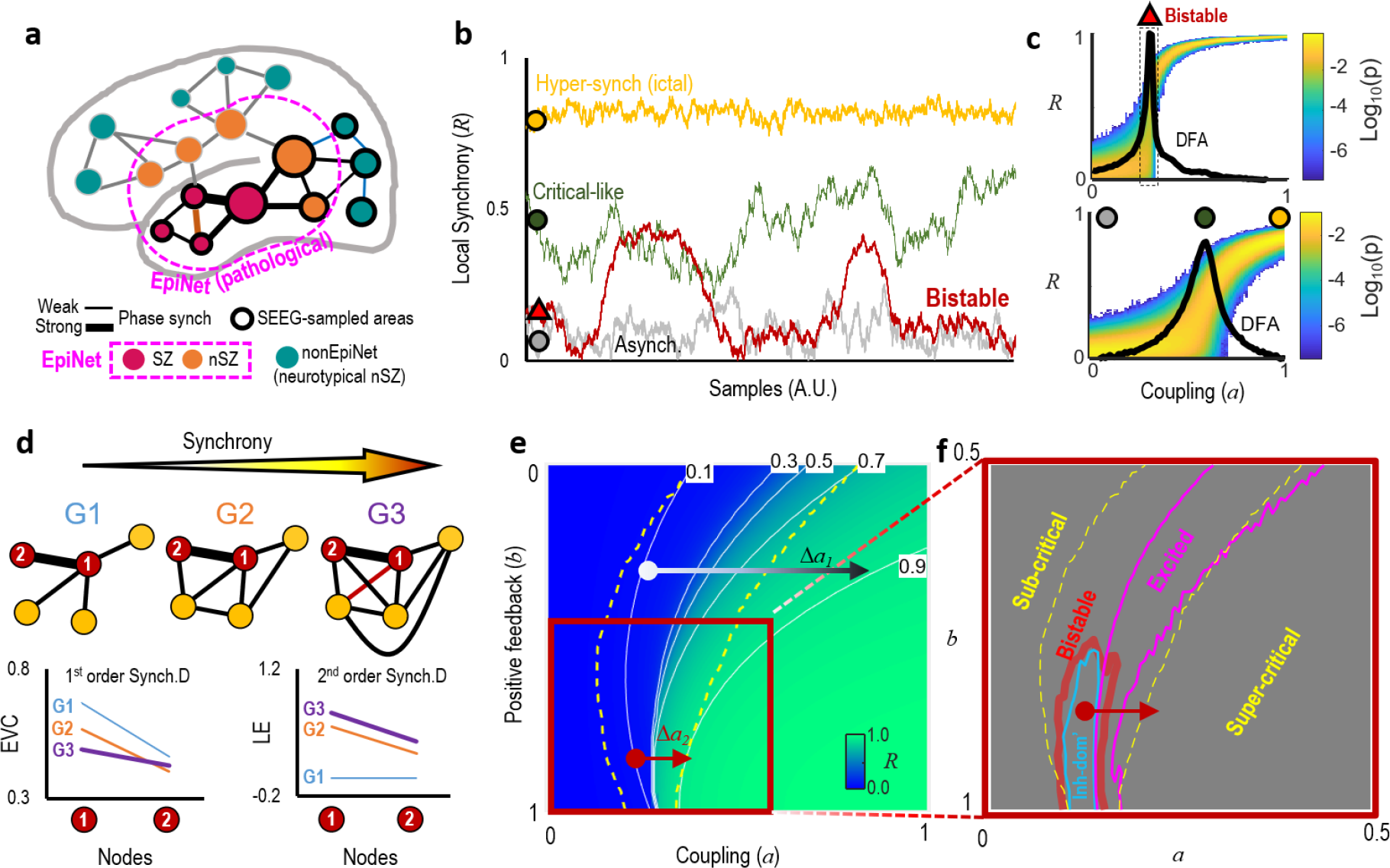
Hypothesis: aberrant local criticality and strong network synchrony concurrently characterize spontaneous activity of the epileptogenic brain. **a.** Seizure-zone (SZ) show seizure activity. The pathological brain network (EpiNet) might comprise the SZ and some non-seizure-zone (nSZ), and the latter were previously considered healthy. SEEG electrode insertion is driven by clinical hypothesis and thus might miss some pathological areas. **b.** Synchrony (*R*), *i.e.*, amplitude envelope of the complex mean-field, when the model is in different regimes. **c.** The probability distribution (*p*) of *R* samples and mean DFA exponents of *R* (black line) as the model was controlled by strong (top) and weak (bottom) local positive feedback; the corresponding time series were in **(b)**; DFA: detrend fluctuation analysis. **d.** As synchrony increased, 1st- and 2nd-order synchrony derivatives of Type 1 (central) nodes are consistently larger than that of Type 2 (peripheral) nodes. The 1st-and 2nd-order refers to a node’s connectivity and its neighbor’s connectivity, respectively. *EV C*: eigen vector centrality; *LE*: local efficiency. **e–f.** Local criticality assessed from spontaneous activity of the model. **e.** Time-averaged *R* as a function of coupling and positive feedback; the area sandwiched between two dashed lines is critical-like regime (*DFA >* 0.6); Δ*a*_1_ and Δ*a*_2_ indicate required increment in coupling to drive the model from critical (R=0.1) to super-critical, seizure-like hypersynchrony (R=0.9) at a given positive feedback strength. The *y* axis is reversed to make this figure comparable with the cusp fold (Supplementary Fig 1). **f.** Association between criticality, bistability, inhibition-dominance, and excitation-dominance in the model local dynamics.

The classic hypothesis of brain criticality postulates that the phase transition is continuous (second-order) [25, 33]. However, both canonical firing rate models [34], ensemble dynamics models [35], and models of synchronization dynamics with positive feedback [36] predict that phase transition may also be discontinuous (first-order) (Fig 1 b). Empirically, operation at a discontinuous phase transition would lead to bistable activity, such as the switching between an UP and an DOWN states during sleep in electrophysiological data from animal models [37, 38] as well as comparable periods of high and low synchrony in awake human resting-state electroencephalography (EEG) [39, 40], magnetoencephalography (MEG), and stereo-EEG (SEEG) [36]. Theoretical work has suggested bistability to occur near criticality when ensembles are influenced by positive feedback [41, 36] or constrained by activity-limiting mechanisms due to over-excitation [42]. Importantly, elevated bistability has been suggested as a universal indicator of a predisposition of the system to exhibit catastrophic forms of aberrant activity [43], such as sudden, uncontrollable bursts of hypersynchrony like those that are observed in various complex systems. We posit here that epileptic seizures could construed as this kind of catastrophic runaway neuronal activity. This notion thus implies that bistability in spontaneous neuronal network activity could be a mechanistic predictive marker for seizure risk [36].

In epilepsy patients’ brain, an extended epileptogenic-like network (**EpiNet**) could entail multiple, possibly overlapping components (Fig 1a), some of which might not engage in every clinically observed seizure or exhibit currently well-known pathological characteristics [44, 45]. We hypothesized that, regardless of seizure types and pathological substrates, regions within the EpiNet would spontaneously operate in a critical regime of high-stability that primes them to catastrophic seizure events. This local aberrant criticality should also be associated a large-scale super-critical trend exhibited as elevated phase synchrony [24] (Fig 1d). Therefore, we evaluated the potential of combining criticality, bistability, and inter-areal synchrony to serve as mechanistic biomarkers.

We tested the hypothesis on retrospective inter-ictal resting-state SEEG from 64 DRE patients. We assessed criticality and synchrony for the SEEG contacts, which were used as features to train supervised classifiers for SZ on cohort level and within subjects. We subsequently used unsupervised classification to investigate whether there were nSZ samples shared similar pathological features with the SZ. This hypothesis-free classification does not aim to classify SZ but to identify an extended pathological brain network, *i.e.*, the EpiNet .

## 2 Materials and Methods

### 2.1 Subjects and SEEG recording

The SEEG were recorded from 64 DRE patients (mean±std age: 29.7±9.5, 29 biological females, Supplementary Table 1) at the Niguarda “Ca’ Granda” Hospital, Milan, Italy [6]. These patients: ***i)***were the first time to undergo SEEG procedure; ***ii)***had no previous brain surgeries, ***iii)***were free of cognitive impairment, psychological, or neurological conditions, and ***iv)***all had been diagnosed with focal onset seizures. Nine subjects who had less than five contacts located in seizure-zone (SZ) or non-seizure-zone (nSZ) were excluded from the SZ-classification analyses. Forty-five subjects underwent surgery or radio-frequency thermo-coagulation (RF-TC) [15] were used for studying the correlation between neuronal features and surgical outcome. In line with [6], a patient’s cumulative surgical outcome with a minimum follow-up of 24 months is considered as “favorable” when meeting the criteria of International League Against Epilepsy (ILAE) classes 1 and 2 (corresponding to Engel classes Ia–Ic). The outcome is considered as “unfavorable” when a patient was not free of disabling seizures (ILAE classes 3–6, Engel classes II–IV).

We obtained 10-min resting-state SEEG as monopolar local-field potentials (LFPs) with shared reference in the white-matter away from the SZ locations [46]. There were no seizures at least one hour before or after the resting-state recording. All patients were under anti-seizure medication with a large variability in the compounds and dosage. The recording time from the last drug administration was not controlled, and therefore the drug effects was not considered.

SEEG leads were platinum-iridium, multi-lead electrodes (DIXI medical, Besancon, France), each of which has 8 to 15 SEEG contacts that were 2 mm long, 0.8 mm thick and had an intercontact border-to-border distance of 1.5 mm. Freesurfer software [47] was used for extracting cortical parcels from pre-surgically obtained T1 MRI 3D-FFE. An automated SEEG contact localization method was then used to assign each SEEG contact to a cortical parcel with submillimeter accuracy (github.com/mnarizzano/DEETO [48]; github.com/mnarizzano/SEEGA [49]).

### 2.2 The seizure-zone (SZ) were clinically identified by physicians

SZ contacts were identified by visual analysis of the SEEG traces. This procedure was carried out by one expert and validated by another [15]. Various peri-ictal and ictal events may initiate from the seizure onset zone (SOZ), including low-voltage fast discharge, spike-and-wave, and poly-spike slow bursts, etc [4]. The brain areas inside the seizure propagation zone (SPZ) do not initiate ictals and often show delayed, rhythmic modulation after seizure initiation from the SOZ [16, 18]. It is also common to see some SEEG contacts to have been identified as SOZ and SPZ simultaneously (SOPZ) by physicians. In this study, we referred to SOZ, SPZ, and SOPZ collectively as the seizure-zone **SZ**.

### 2.3 Preprocessing and filtering

Contacts located in SZ are known to demonstrate inter-ictal events (IIE) characterized by high-amplitude spikes with widespread spatial diffusion, which could bias criticality estimates. We followed the approach used in [50] to identify and exclude IIEs from biasing the assessments. Briefly, each contact broad-band signal was first partitioned into non-overlapping 500 ms segments; a segment was tagged as ‘spiky’ and discarded from LRTCs and bistability analyses when at least 3 consecutive samples exceeding 7 times the standard deviation above the channel mean amplitude. Last, narrow-band frequency amplitude time series was obtained by convoluting the broad-band SEEG contact time series with Morlet wavelets (m=5) from 2 to 225 Hz with equal inter-frequency distance on *log*_10_ scale.

### 2.4 Assessing long-range temporal correlations (LRTCs)

After filtering, continuous narrow-band neuronal oscillations were subject to criticality assessments (the formal definitions of the metrics and exemplary SEEG traces can be found in the Supplementary Methods).

The critical exponent obtained using linear detrend fluctuation analysis (DFA) is conventionally used to assess the LRTCs in the ongoing narrow-band oscillation [27]. The DFA exponent quantifies how the root mean square of local fluctuations grows with logarithmically increasing sampling window size. In other words, the exponent could predict how fast the fluctuations would grow in the long-run with only a small fraction of all available data. *DFA* = 0.5 indicates that the time series is indistinguishable from a random walk process with no long-range memory; 0.5 *< DFA <* 1 indicates significant LRTCs, *i.e.*, an indication of criticality; 1 *< DFA <* 2 indicates the time series is non-stationary.

### 2.5 Assessing bistability

The BiS index quantifies the degree of bistability of a neuronal oscillation [39]. First, the empirically observed probability distribution of the narrow-band power time series is fitted with a single exponential and a bi-exponential model. Then, the BiS is derived from the model comparison between the two competing models based on Bayesian information criterion of the two candidate models. A *BiS >* 3 indicates bistability.

### 2.6 Assessing functional excitation/inhibition (E/I)

The functional excitation/inhibition (E/I) index (*f E/I*) is closely related to the DFA. It makes inferences about the operational regime of the E/I population based the hypothetical relationship between time-resolved scaling exponent and local synchrony [29]. There could be three underlying operational regimes: 0 < *f E/I* < 1 indicates that the oscillation demonstrated inhibition-dominated temporal dynamics; 1 < *f E/I* < 2 indicates excitation-dominated dynamics; and *f E/I* = 1 with a constraint of 0.5 < *DFA* < 1 indicates an excitation-inhibition balanced dynamics.

### 2.7 Assessing and validating phase synchrony derivatives

To examine whether large-scale brain networks were characterized by hypersynchrony associated with a trend towards super-criticality within the critical regime, we conducted graph analyses on the all-to-all phase synchrony between SEEG contacts, *i.e.,* connectomes. A connectome was treated as a *graph*, wherein each contacts are *nodes* and inter-contact synchrony are *edges* [51]. If the EpiNet indeed shows hypersynchrony, the nodes in the EpiNet would be characterized by elevated 1st- and 2nd-order centrality (Fig 1d) [52], *i.e.*, a node’s own and its neighbors’ connectivity, respectively.

The synchrony between all contacts located in the cortical and subcortical gray matter was assessed using the phase-locking value (PLV) for 50 narrow-band frequencies (Morlet wavelets, *m* = 7.5). The frequency bands spanned from 2 to 450 Hz with equal inter-frequency distance on *log*_10_ scale. Across subjects, a small number of contacts were referenced with the same white-matter contact. The PLV edges between them were set to zeros as recommended by[46], *e.g.*, Fig 2f.

**Figure 2.**
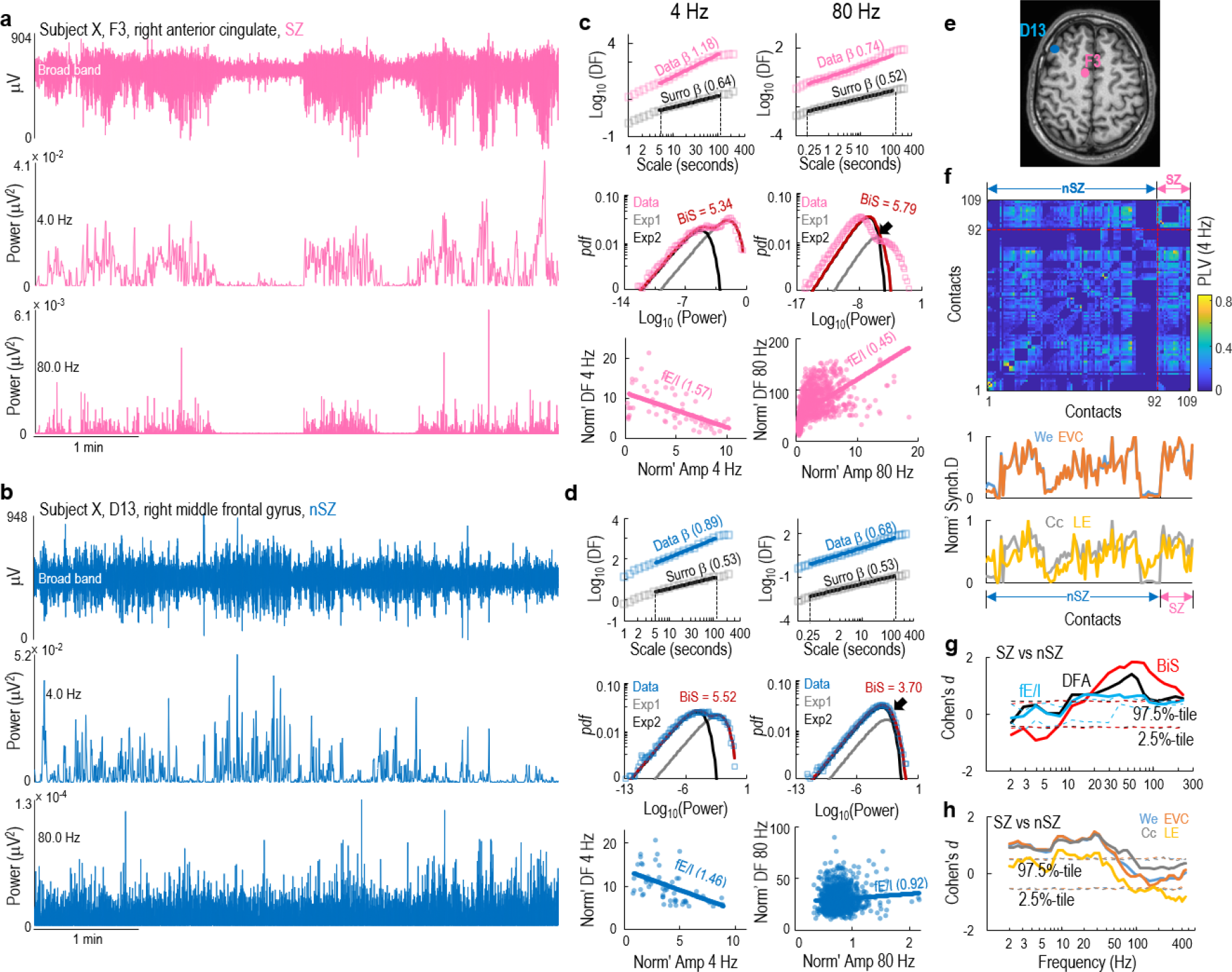
Individual level evidence of differences between SZ and nSZ. **a–b**. Five minutes of broad-band and narrow-band traces from **(a)** a SZ contact and **(b)** a nSZ contact from frontal region of subject *X*. **c.** The fitting of DFA exponent (top), the BiS (middle), and fE/I (bottom) for 4 Hz and 80 Hz oscillation from **(a)**. Markers and solid lines indicate observed data and fitted models, respectively. DF: detrend fluctuation; *pdf* : probability distribution function; Norm’: normalized; Amp: amplitude envelope. **d.** The same as **(c)** for **(b). e.** SEEG contact locations for **(a)** and **(b). f.** Top: subject *X*’s 4 Hz phase synchrony between all contacts assessed using phase-locking value (PLV). Bottom: normalized synchrony derivatives of the SEEG contacts from the PLV matrix. We: effective weight; EVC: eigen vector centrality; Cc: clustering coefficient; LE: local efficiency. **g–h** Difference between SZ and nSZ in **(g)** narrow+band criticality assessments and **(h)** synchrony derivatives. Dashed line indicate confidence intervals observed from 1,000 label shuffled surrogate data.

To characterize SEEG contact connectivity, the PLV connectomes were collapsing into 1D synchrony derivatives [52] using the Brain Connectivity toolbox (brain-connectivity-toolbox.net). We estimated 1st-order synchrony derivatives, *i.e.*, the connectivity of a given contact, with *effective weight* (*We*) and *eigenvector centrality* (*EV C*). *We* is the linear aggregating available edges. *EV C* is a self-referential measure of centrality, *e.g.*, nodes have high *EV C* if they connect to other nodes that have high *EV C*. We estimated 2nd-order synchrony derivatives, *i.e.*, the connectivity of the neighbors of a given contact, with *clustering coefficient* (*Cc*) and *local efficiency* (*LE*). *Cc* is the fraction of node’s neighbors that are neighbors of each other. The *efficiency* is defined as the average inverse shortest path length in the network, and *LE* is the *efficiency* assessed in the neighborhood of a node *n*_*i*_, *i.e.*, all the nodes connected to *n*_*i*_.

Spatial sampling inhomogeneity of SEEG could be exacerbated by the deletion of PLV edges due to shared-reference, which might bias the properties of the synchrony graphs. Before analyses, we ascertained that the synchrony derivatives used here were robust against missing samples by exhaustive simulations. This was tested with two sets of simulations. First, for each subject’s narrow-band PLV matrices, random edge deletion was used to emulate the removal of interactions between contacts that shared the same white-matter contacts. Second, node deletion was used to emulate the individual variability in spatial sampling and therefore possible sub-sampling. All these synchrony derivatives were consistently robust against simulated random deletion (Supplementary Fig 4).

### 2.8 Normalizing, frequency clustering, and differentiating brain dynamics features

We first differentiated SZ and nSZ contacts within subjects and then aimed to train cohort-level models for supervised SZ-classification. The criticality and phase synchrony estimates were known to show co-variability and considerably large individual variability across subjects [24]. Here, we concerned the difference between neurotypical and pathology within subjects, and therefore, the criticality and synchrony derivatives were normalized as: 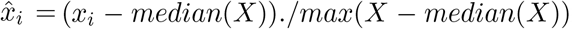, where, *x*_*i*_ is a contact, *X* is a 1D vector of real numbers of criticality or synchrony derivative for all contacts for a given frequency within a subject (Fig 3 a–f).

**Figure 3.**
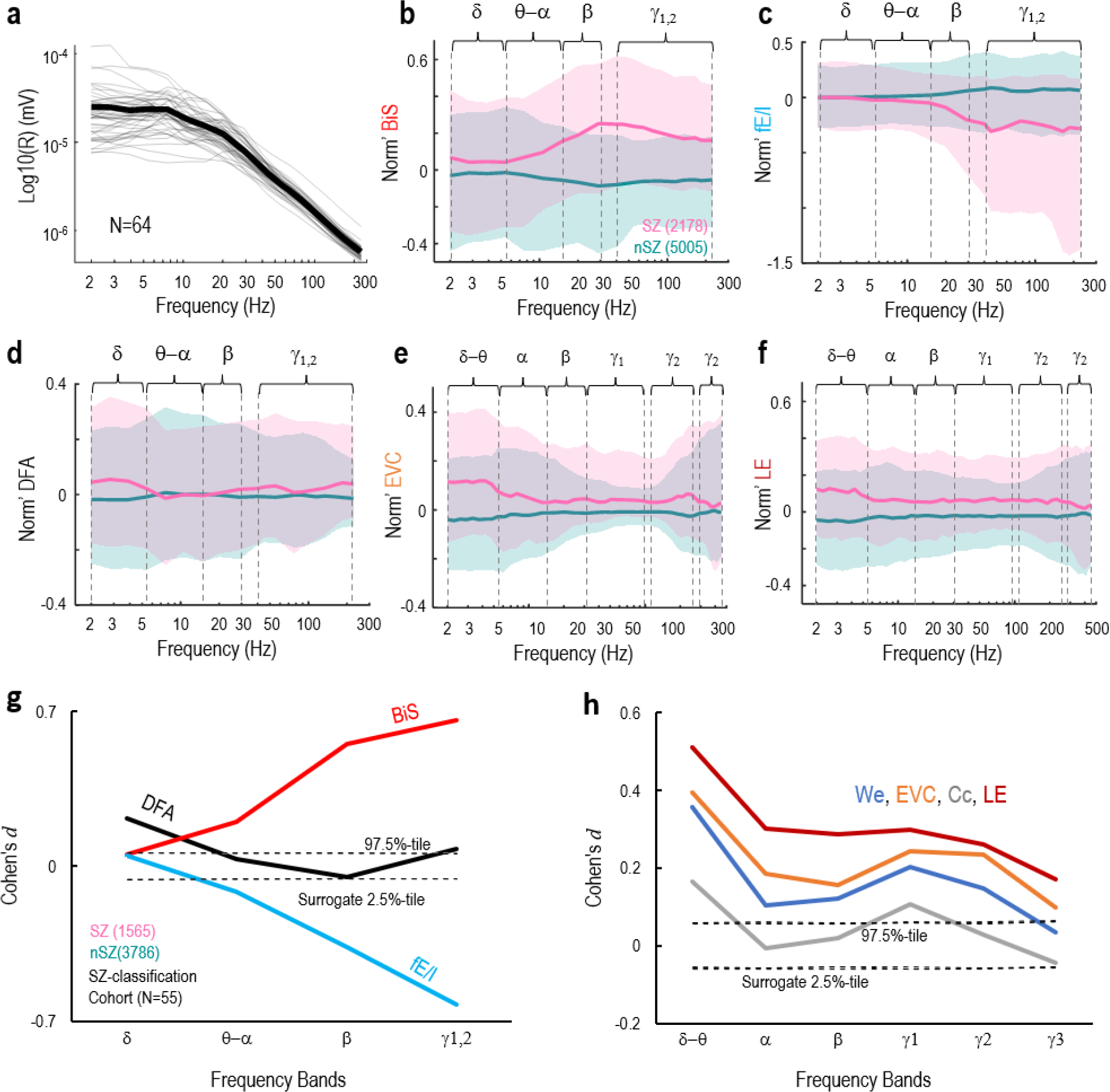
The EZ showed concurrent anomalies in criticality and synchrony assessments. **a–f**. Demographics of the whole cohort samples (N=64). **a.** With-subject (thin) mean SEEG amplitude and group average (thick). **b–f.** Normalized narrow-band frequency criticality and synchrony derivatives for SZ and nSZ contacts. BiS: bistability index; fE/I: functional E/I index; DFA: detrend fluctuation analysis exponent; EVC: eigen vector centrality; LE: local efficiency; Shades indicate 25 and 75%-tile; frequency clusters of criticality: *δ* (2–4 Hz), *θ* −*α* (5.4–11 Hz), *β* (15–30 Hz), and *γ*_1,2_ (45–225 Hz); frequency clusters of synchrony derivatives: *δ* −*θ* (2–5.4Hz), *α* (6.1–13 Hz), *β* (15–30 Hz), *γ*_1_ (40–96 Hz), *γ*_2_ (110–250 Hz), and *γ*_3_ (270–450 Hz). **g–h.** Differences between SZ and nSZ in band clustered metrics for the sub-cohort (N=55)used in supervised SZ-classification. Dished lines indicate confidence interval observed from 10^4^ label-shuffled surrogates conducted independently for each metric.

Because the topological features of narrow-band criticality assessments were similar between neighboring frequencies and different between fast and slow brain rhythms, twenty narrow-band criticality assessments were next collapsed into four frequency clusters based on observed similarity as *δ* (2–4 Hz), *θ* −*α* (5.4–11 Hz), *β* (15–30 Hz), and *γ*_1,2_ (45–225 Hz) (Supplementary Fig 6); Likewise, as we previously reported that the narrow-band *PLV* matrices showed topological similarity [50], fifty narrow-band synchrony derivatives were collapsed into six frequency clusters as: *δ* −*θ* (2–5.4Hz), *α* (6.1–13 Hz), *β* (15–30 Hz), *γ*_1_ (40–96 Hz), *γ*_2_ (110–250 Hz), and *γ*_3_ (270–450 Hz) ([50]).

Subsequently, the effect size of differences between SZ and nSZ in these normalized and frequency clustered feature data were assessed with Cohen’s *d* and compared with the 99%-tile of Cohen’s d observed from 1,000 label-shuffled surrogates (Fig 3 g–h).

### 2.9 Supervised SZ-classification using the Random Forest algorithm

We first assessed the individual feature importance and then conducted supervised SZ-classification using the SZ labels identified by physicians. The feature importance of these neuronal estimates were assessed with the SHapley Additive exPlanations (SHAP) values [53]. The SHAP values is a generic metric to explain any tree-based model by explicating the local and global interpretability of features, which advances the transparency that conventional “black-box” classifications approaches lack of. The non-parametric Random Forest algorithm was employed for supervised SZ-classification [54]. The algorithm is a machine learning method uses bootstrapped training data and combines the simplicity of decision trees with extended flexibility to handle new data.

### 2.10 Unsupervised classification by contact cluster analysis

We also conducted unsupervised contact classification by cohort level contact cluster analysis, *i.e.*, without considering any human inputs as the ground truth for pathophysiology. This was done by first pooling all feature data of the whole SZ-classification cohort (contacts *×* features, 2D scalars), then computing all-to-all feature-similarity (*i.e.*, Spearman’s rank correlations *r*) between contacts (Fig4e), and last conducting clustering analyses using the unweighted pair group method with arithmetic mean (UPGMA) [55]. The UPGMA is an agglomerative hierarchical clustering approach that builds a hierarchical tree through an iterative procedure to reflect the distance between all pairs of objects (*i.e.*, either a contact or a cluster of contacts) represented by the similarity matrix. The information about distance tree structure is then used to partition the data into well delineated clusters.

**Figure 4.**
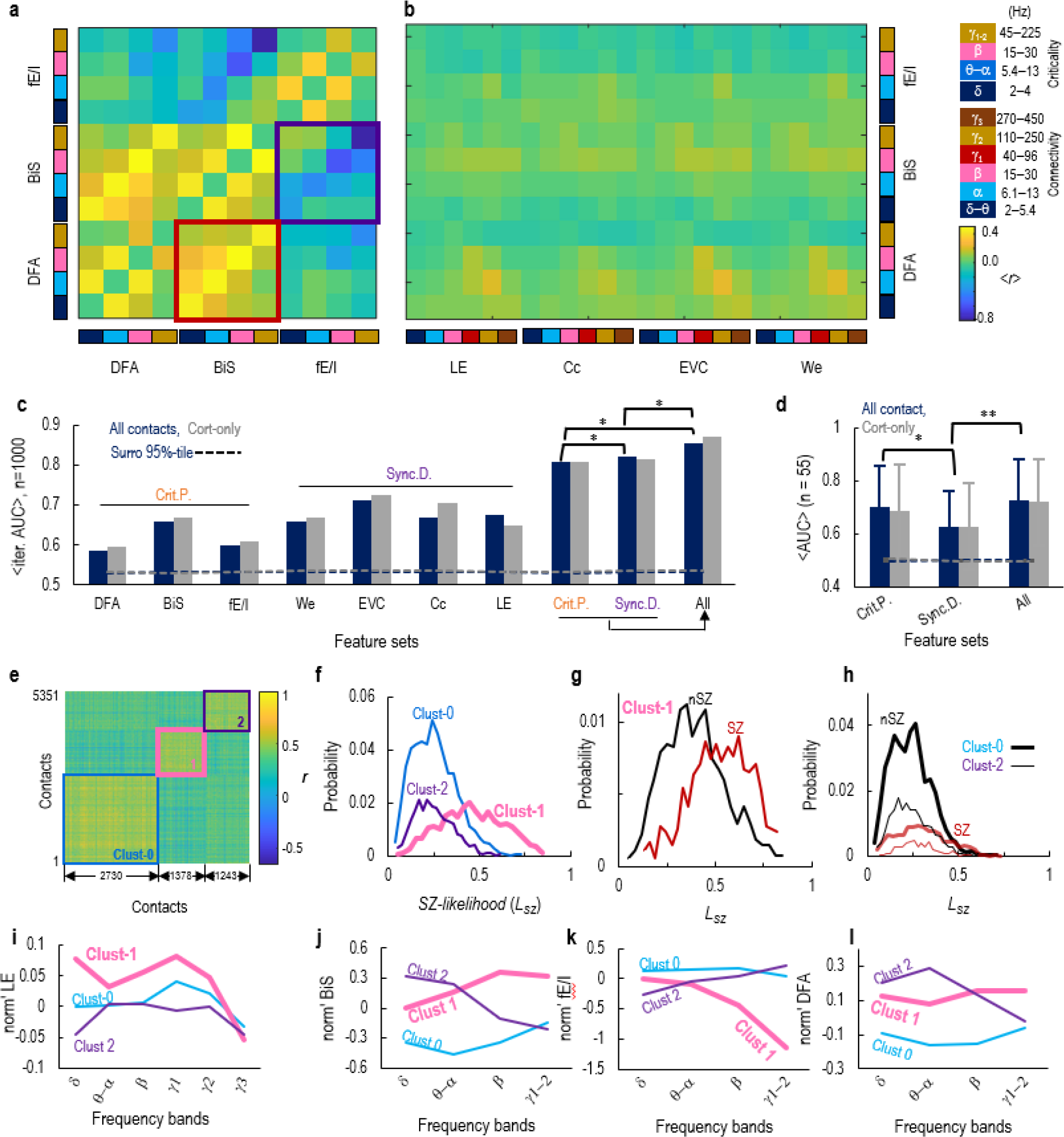
Optimal SZ-classification was achieved by combining all criticality and synchrony features. **a.** Mean within -patient Spearman’s correlation (*< r >*) between band-collapsed criticality assessments. **b.** Similarity between criticality and connectivity across frequencies. **c.** Population pooled SEEG contacts classification cross-validation (randomly split 20% vs 80% test vs train set over 1000 iterations) when using different feature set. AUC: mean area under the receiver-operator-curve; * indicates differences (repeated t-test, −*log*10(*p*) in sequence: 62.1, 39.9, 117.8, Benjamini-Hochberg FDR corrected); dashed line: surrogates mean (*n*_*surro*_= 1000). **d.** Mean subject AUC (n=55) of within-patient SZ-classification; * and **: indicate differences (unpaired t-test, −*log*10(*p*) = 2.2 and 6.1, respectively (FDR corrected); error bars: standard deviation; dashed lines: surrogate mean (*n*_*surro*_= 1000). **e.** All-to-all SEEG contact similarity matrix, and the boxes demarcate three clusters. **f.** Distribution of SZ-likelihood for each cluster defined in (**e**). **g–h.** *L*_*SZ*_of SZ and nSZ of (**g**) cluster-1, (**h**) cluster-0 and cluster-2. **i–l.** Intra-cluster group average of (**i**) LE, (**j**) BiS, (**k**) fE/I, and (**l**) DFA. Inter-cluster differences in theses features see Supplementary Fig 8 (e–f).

## 3 Results

### 3.1 Positive feedback in a computational model of synchronization dynamics leads to bistability and increased risk for seizure-like hypersynchrony

To investigate the emergence of seizure-like hypersynchrony in systems with separately controllable bistability and criticality, we used a Kuramoto model [24, 56] equipped with positive feedback [36]. The Kuramoto model is a generative model for synchronization dynamics, *i.e.*, a model that generates time series of oscillations characterized by emergent synchronization that is quantified by “order” (*R*) so that *R* = 0 in the absence of synchrony and *R* = 1 for perfect synchrony (Supplementary eq.6). The positive feedback is a generative mechanism for bistability in canonical systems such as the cusp bifurcation that also considers two control parameters [43]. Based on the cusp bifurcation, we expected: *i)* the model order to be influenced by both coupling (*a*) and positive feedback (*b*); *ii)* bistability in order should emerge exclusively with increasing positive feedback ([36], Supplementary Fig 1).

At low levels of feedback, increasing coupling led to emergence of power-law scaling LRTCs at moderate levels of order – indicating a smooth phase transition in the order consistent with the prediction of the classic criticality hypothesis (Fig 1e, see also Supplementary Fig 2). However, in the presence of stronger feedback, the model exhibited bistable synchronization dynamics within the critical regime (Fig 1f).

Importantly, in the regime of critical bistability, the model became hyper-sensitive to changes in coupling and could erratically transit from criticality to super-criticality. For example, a 15% increase in coupling could drive the model from the subcritical side of the critical regime with low synchrony (*R* = 0.1) into a super-critical regime with seizure-like hypersynchrony (*R* = 0.9); as a contrast, at low levels of feedback, a 61% increase in coupling was required to drive the model to exhibit the same subcritical-to-supercritical transition (see Δ*a*_1_ vs Δ*a*_2_ in Fig 1e).

Lastly, within the regime of critical bistability, the excitation-dominant dynamical regime is closer to the super-criticality than the inhibition-dominant dynamical regime (Fig 1e, Supplementary 2).

### 3.2 Seizure-zone (SZ) exhibits aberrant criticality and elevated synchrony

We assessed local criticality and inter-areal phase synchrony in inter-ictal, resting-state SEEG recordings of local neocortical field potentials, and then examined the capacity of these synchronization dynamics metrics to differentiate the SZ from non-seizure-zone (nSZ).

The amplitude envelope of neuronal oscillations reflects local synchronization and is equivalent to the model order, and local criticality of the narrow-band SEEG data was assessed in the same manner as for the model. We assessed all-to-all phase synchrony between all SEEG contacts with the phase locking value (*PLV*), which was then used to obtain the first- and second-order synchrony derivatives, *i.e.*, a contact’s own and its neighbors’ connectivity, respectively. (Fig 1d).

Visual inspection of individual data revealed differences between SEEG contacts located in SZ and nSZ. SZ contacts often exhibited stronger bistability in broad-band and narrow-band traces than adjacent nSZ contacts (Fig2 **a–b**, Supplementary Fig 3). In this representative subject, the SZ was characterized by more pronounced bistability, stronger LRTCs, and more inhibition-dominance in the 80 Hz than the nSZ (Fig2 **c–d**). Moreover, the 4 HZ PLV connectome showed that both the synchrony among SZ contacts (first-order connectivity) and between the SZ and nSZ contacts(second-order connectivity) were pronounced (Fig2 **e–f**). The differences between SZ and nSZ in narrow-band criticality and synchrony assessments in this subject are shown in (Fig2 **g–h**).

For the whole cohort, LRTCs, bistability, and functional E/I were assessed for 2 – 225 Hz narrow-band oscillation amplitudes (Fig 3**a**). Pooling contacts from all subjects revealed that the SZ appeared to have elevated bistability and inhibition-dominance in 45–225 Hz and 15–30 Hz (Fig 3**b, c**), whereas slightly stronger LRTCs in 2–5 Hz and above 100 Hz (Fig 3**d**). Meanwhile, the SZ showed elevated first- and second-order synchrony derivatives (Fig 3**e, f**, respectively). Overall, these narrow-band criticality and synchrony measures exhibited highly similar anatomical patterns between neighboring bands but were less similar between distinct frequency bands (Supplementary Fig 6). To improve interpretability and reduce dimensionality, we collapsed the individual frequencies into bands and used them to assess the statistical differences between SZ and nSZ. For criticality, the 20 narrow-band maps were collapsed into four bands (see Fig 3**b–d** and Supplementary Fig 6). Similarly, the 50 narrowband-synchrony-derivative maps were collapsed into six bands (see Fig 3**e, f**).

Comparing to the nSZ, the SZ exhibited greater bistability with concurrent stronger inhibition in *γ*_1,2_ (40 − 225 Hz) and *β* (15 − 30 Hz) band oscillations (Fig 3**g**). This form of aberrant criticality was highly similar to the model dynamics in the high seizure-risk regime. Furthermore, the SZ showed simultaneously elevated first- and second-order synchrony derivatives in *δ* −*θ* (2 − 5.4 Hz) followed by *γ*_1_ (40 − 96 Hz) band (Fig 3**h**). This indicated that SZ were central nodes (*i.e.*, the Type-1 node, Fig 1**d**) characterized by elevated resting connectivity to SZ and between the neighbours of SZ. Finally, these findings indicate that the inter-ictal brain dynamics in SZ are statistically dissociable from those in nSZ in terms of aberrant bistability, classical criticality, and excitability.

### 3.3 Supervised SZ classification: combining criticality and synchrony assessments maximizes SZ-classification accuracy

Next we asked if the criticality and synchrony derivatives could be used to classify the SZ identified by physicians, which constitutes the first prerequisite for this approach to have clinical value. To this end, we used a supervised learning approach and trained models to identify the SZ contacts in individual patients in three steps. First, we assessed the similarity between criticality and synchrony derivatives. Next, as a proof-of-concept, we conducted a population level cross-validation. We employed the Random Forest algorithm that is a supervised Bayesian classifier [54], and the SZ contacts were used as the ground truth for training. Lastly, we conducted within-patient leave-one-out SZ-classification, *i.e.*, one subject as the test set and the rest as the train set.

First, feature similarity was assessed as correlations between band-collapsed local criticality and synchrony features (Supplementary methods). The bistability index (BiS) and DFA were positively correlated between all bands (Fig 4**a**, respectively). In line with the modeling results, the BiS and fE/I indices were negatively correlated which constituted the first empirical evidence of concurring bistability and inhibition-dominance in SZ to support the model prediction of elevated risk – but not immediately prior to supercriticality, *i.e.*, a high-bistability and excitation-dominance regime (Fig 1**e–f**). The weak correlation between criticality and synchrony derivatives (Fig 4**b**) implied that they reflected non-overlapping physiological processes. Since both feature families showed differentiating effects for SZ (Fig 3), combining them would most likely offer better SZ-classification than using any individual features alone.

Next, as a precaution, we took two additional procedures to validate that criticality and synchrony assessments were useful features for SZ-classification. First, we assessed global and within-subject feature importance with the Shapley Additive exPlanations (SHAP) values for the Random Forest classifier [53]. This revealed that, on the population level, *γ* band BiS and fE/I, *β* band BiS, and *δ* −*θ* band LE were among the most important individual features (Supplementary Fig 7). Next, we conducted a population level cross-validation with 1, 000 independent iterations, each of which was a random 80 : 20%-partition (training:test set) (Supplementary methods). We tested SZ-classification using criticality alone, synchrony derivatives alone, and combining criticality and connectivity and for “cortical contacts only” and “all contacts” (*i.e.*, cortical and subcortical). These tests revealed that combining criticality and connectivity (‘All’ in Fig 4c) yielded the best classification accuracy with the area under curve (AUC) of the receiver operating characteristic reaching 0.85 *±* 0.002 (mean *±* std). These results thus offered proof-of-concept for within-patient SZ-classification.

Lastly, we performed within-patient SZ-classification with leave-one-out validation, wherein each patient’s contacts served as the test set (n=55, patients with less than five SZ or nSZ contacts were excluded). Using all features yielded the best result with a mean AUC of 0.73*±*0.16 for all contacts, and 0.72*±*0.16 for cortical contacts only (Fig 4 **d**), with no difference between cortical-only and all contacts (repeated t-test, *p >* 0.55). Synchrony derivatives were more potent than criticality for SZ-classification in population cross-validation, whereas criticality represented more potent features than connectivity in within-patients classification.

### 3.4 Unsupervised classification reveals a pathological EpiNet sample cluster

The SZ and nSZ exhibited distinct brain dynamics, and these differences could be leveraged to classify SZ and nSZ in individual patients with moderate-to-high accuracy. We next asked whether the pathological brain network, the EpiNet, could entail not only the SZ but also brain areas clinically identified as healthy nSZ. To answer this question, we used unsupervised classification. We first estimated all-to-all feature similarity between contacts from all subjects and then conducted clustering analyses of this cohort level inter-contact similarity matrix (Supplementary methods). Thereby, contacts belonging to the same cluster are more similar with each other than with the contacts from other clusters. We found that through a range of partitioning resolutions from two to nine clusters, three major clusters remained stable in their constituent contacts and were representative of the whole cohort (Supplementary Fig 8 **a–b**). Thus, we chose the 3-cluster partition solution for further analyses (cluster size *n* = 2730, 1378, 1243, respectively) (Fig 4e).

We subsequently checked if the clusters represent distinct (patho-)physiological profiles by joint analyses of the clustering and within-patient supervised SZ-classification. Based on the similarity between test set and the SZ in the train data, the supervised classifier assigned a SZ-likelihood (*L*_*SZ*_, 0 − 100%) to each contact in test set to indicate how likely it was a SZ. The Cluster-1 contacts (pink, Fig4 **f**) showed larger mean *L*_*SZ*_ than Cluster-0 and -2 (unpaired t-test −*log*_10_(*p*) *>* 10^15^ and *>* 234.2, respectively), whereas there was no difference between cluster-0 and -2 (unpaired t-test, −*log*_10_(*p*) *<* 0.35). Here, a larger *L*_*SZ*_ indicated that pathological-like contacts were indeed more concentrated in Cluster-1 (in-cluster SZ:nSZ = 48.8% : 51.2%). The probability of observing SZ in Cluster-1 (48.8%) were more than twice larger than that of Cluster-0 and -2 (23.0% and 21.2%, respectively), which further supported that Cluster-1 represented pathophysiology, *i.e.,* the hypothetical EpiNet.

We next asked what features differentiated the three clusters by comparing criticality and synchrony assessments between clusters. Cluster-1 showed elevated local efficiency (Fig 4**i**) and eigen vector centrality (Supplementary Fig 8f) most prominently in *δ* −*θ* and *γ*_1_ band. The concurrent larger first- and second-order synchrony derivatives suggested that these brain areas were relatively central (Type-1 node, Fig 1**f**). Cluster-1 also showed elevated bistability (Fig 4**j**) with concurring inhibition-dominance (Fig 4**k**) in *γ*_1,2_ and *β* band comparing to Cluster-0 and -2. Lastly, Cluster-1 showed higher DFA in *γ*_1,2_ band (Fig 4**l**).

Notably, in the tentatively pathological Cluster-1 for EpiNet, 25.2% of the nSZ contacts were identified as SZ by the supervised classifier (*L*_*SZ*_ > 50%, Fig 4g). Hence, the the ratio of such ‘pathological-like’ nSZ contacts (*p*_*nSZ*_−*clust*1) in a patient might tell the extent of the EZ network that could not be detected using conventional approaches. The *p*_*nSZ*_−*clust*1 was higher in patients with predominantly frontal SZ (4.7 *±* 3.8%, *n* = 15) than the patients with SZ in temporal or other lobes (2.6 *±* 2.6%, *n* = 40) (unpaired t-test, *p <* 0.018), whereas the mean *L*_*SZ*_ of all nSZ contacts in the frontal-SZ patients (23.6 *±* 4.9%, *n* = 15) and other patients (24.5 *±* 7.1%, *n* = 40) had no difference (unpaired t-test, *p >* 0.667). This suggested that these tentatively pathological nSZ were undetected during pre-surgical EZ mapping in the frontal lobe seizure patients, who have previously been suggested to likely represent a special type of focal epilepsy [57].

## 4 Discussion

With our putative mechanistic biomarkers for epileptogenicity, we extracted neuronal features from inter-ictal SEEG and trained supervised classifiers to localize the SZ against gold-standard clinical localization. The employed novel biomarkers were derived by using computational modeling motivated by complex system, brain criticality, and catastrophe theory. They include bistability, long-range temporal correlations, and functional excitation/inhibition balance in local synchrony dynamics. In the model, strong positive feedback led to a high seizure-risk regime, wherein the local synchrony exhibited high bistability, strong inhibition, and aberrant power-low scaling. The SEEG analyses revealed that the local synchrony in SZ areas exhibited striking similarity to the model in the high risk regime. Meanwhile, the SZ areas simultaneously exhibited strong inter-areal synchrony, implying large-scale super-criticality. Supervised classifier trained on all criticality and synchrony features yielded more accurate (AUC 0.85) identification of SZ than using any individual feature alone (AUC 0.6–0.7).

We subsequently investigated whether the brain network exhibiting pathophysiological features constitutes more extended areas (EpiNet) than the clinically localized SZ. Unsupervised classifiers yielded three principal sample clusters, among which only one was pathological-like. Co-localization of the supervised and unsupervised approaches further showed that the pathological cluster included electrode contacts with the greatest likelihood of being SZ. Importantly, however, 50% of contacts in the pathological cluster were considered healthy in clinical assessment.

### 4.1 Strong positive feedback leads to elevated bistability, strong inhibition, and lowered hypersynchrony threshold

Positive feedback has been suggested to cause bistability in complex systems irrespective of their physical details [41]. Excessive bistability indicates a first-order discontinuity with harmful hysteresis [36] and phenomenologically a signature of incoming catastrophes [43, 58]. In both unimodal and bistable critical regimes, increasing coupling strengths can lead to a seizure-like hypersynchronous regime. However, only the bistability regime is associated with a lowered threshold to hypersynchrony, wherein a small increase in coupling can result in a sudden regime shift from asynchrony to full synchrony. Within the model critical bistability regime, the excitation(E)-dominant dynamical regime was closer to the supercriticality than the inhibition(I)-dominant dynamical regime. In the SEEG, the SZ exhibited larger bistability and predominantly I-dominance, rather than E-dominance, in *β* −*γ* bands. These novel results thus support the cusp catastrophe prediction and further suggest that concurrent high bistability and strong inhibition – in a critical like regime – characterize an increased seizure risk.

Theoretical work has shown that positive feedback could be associated with high resource demand with ensuing bistable neuronal avalanche dynamics [42]. Brain regions with high resource costs are considered vulnerable to abnormal development and pathology that lead to disorders [59]. In our simple model, strong positive feedback was operationalized as a state-dependent term. In more realistic models such as the Wilson-Cowan ensemble, several synaptic mechanisms including strong E-to-E self-excitation and E-to-I disinhibition can lead to bistable firing rates [34, 60]. These arguments have also been further supported by recent evidence on the positive feedback between seizure and metabolic anomalies [61, 62] and neuroinflammation [63, 64].

### 4.2 The EZ is a hypothetical core of the epileptogenic network

The concept of EZ has been evolving over the decades [44, 16, 45]. Nonetheless, when defining the EZ, the clinical objective of achieving long-term seizure freedom has remained unchanged. [65]. As the EZ represents the core of the epileptogenic network to generate seizures, and hypothetically, there could be several possible surgical solutions to be weighted by clinical constraints [12].

Across individuals, the EZ [16, 18, 66] might consist components such as structural lesions, seizure zone, high gamma oscillators, and irritative zone, etc [45, 44], some of which may overlap. This could account for why individual markers often localize EZ inconsistently across subjects [8, 9]. The low post-surgical seizure freedom in many patients has further suggested that the pathological networks were not fully mapped with existing conventional biomarkers. We found that combining all criticality and synchrony biomarkers yielded the highest classification accuracy compared to using any single biomarkers alone. This finding thus offered evidence to support the multi-component and our concurrent local and global pathology hypotheses.

### 4.3 The EpiNet represents a pathological brain network that is more extended than previously thought

With unsupervised classification we revealed an EpiNet cluster entailing both SZ and nSZ with shared pathological characteristics. In these patients, the nSZ did not show seizure activity throughout the SEEG monitoring period spanning from days to weeks and therefore were deemed as non-pathological. However, unsupervised classifiers trained on our novel biomarkers using 10-min of resting-state SEEG could readily reveal that these nSZ shared similar functional profiles with the SZ. These finding suggested that the entirety of the EZ network might indeed encompass brain areas that do not engage in every observed seizure. Alternatively, while not involved in seizure activity, these nSZ might be linked to the altered structural connectivity within the SZ, potentially suggesting regions of future ictal network spread or kindling.

In contrast, some SZ were assigned to the two remaining clusters that did not show clear signs of pathophysiology and were assigned low SZ-likelihood by the supervised classifier. We postulated these atypical SZ might be non-central nodes in the EZ (*i.e.*, Type-2 nodes in Fig 1**d**). Due to the unavailability of the structural connectome data, we could not simulate large-scale synchrony to test this idea. Future efforts should be directed into elucidating the interesting dissociation reported here, in particular the functional role of the nSZ in the EpiNet cluster [45].

### 4.4 Integrating brain criticality and synchrony: novel mechanistic biomarkers for epileptogenicity

The joint consideration of these synchrony and criticality measures opens new avenues for data-driven and automated identification of putative EpiNet areas in individuals, which may complement current clinical tools and improve the outcome of epilepsy surgeries. In this study, physicians identified the SZ case by case based on the SEEG readouts during inpatient monitoring. We used the SZ as the ground truth for training classifier to identify pathological areas. The supervised classification results were subsequently complemented by hypothesis-free unsupervised classification. However, the SZ identification accuracy could be variable across subjects because the spatio-temporal signatures of ictals are complex across seizure types[4], sometimes can be highly ambiguous [67], and dependent on multiple seizure cycles [68, 69]. The EZ, on the other hand, represents a surgical solution for attainment of seizure freedom, and the overlap between the EZ and the EpiNet revealed here is still unclear. The accurate EZ-localization will involve rigorous hypothesis testing and validation, *i.e.*, using virtual surgery and real surgery, respectively. [70, 71].

## 5 Conclusion

We showed that our novel complex-system and systems-neuroscience driven biomarkers were able to discover epileptogenic pathology that were not previously known. Combining these potent biomarkers with state-of-the-art learning approaches offers a promising avenue for comprehensive localization of the epileptogenic network.

## Supporting information

Subject information.

## Author contributions

Conceptualization: SHW, JMP

Funding acquisition: JMP, SP, SHW

Methodology: SHW

Software: SHW, VM, GA

Formal analysis: SHW

Resources: GA, LN

Data Curation: GA, VM

Visualization: SHW, PC, JMP

Writing - Original Draft: SHW, PF, LN, PC, SP, and JMP

Supervision: JMP

## Acknowledgments

This project was funded by: the Academy of Finland grant (SA 253130 and 296304) awarded to JMP; Sigrid Jusélius Foundation grant awarded to SP and JMP; NEXTGENERATIONEU (NGEU) and by the Ministry of University and Research (MUR), National Recovery and Resilience Plan (NRRP), project MNESYS (PE0000006) – A Multiscale integrated approach to the study of the nervous system in health and disease (DN. 1553 11.10.2022) awarded to LN. The theoretical work was partially funded the Ella and Georg Ehrnrooth Foundation awarded to SHW (14-10553-2); the analytical tools development, data analyses, and publication preparation were supported by the Finnish Cultural Foundation postdoc fellowship (00220071) and the Sigrid Jusélius Foundation fellowship (210527) awarded to SHW. SHW is also supported by the DARLING project (Projet-ANR-19-CE48-0002) granted to PC. We are grateful for the support.

## Study Approval Statement

This study was approved by the ethical committee (ID 939) of the Niguarda Hospital, Milan, Italy, and was performed according to the Declaration of Helsinki. We confirm that we have read the Journal’s position on issues involved in ethical publication and affirm that this report is consistent with those guidelines.

## Patient Consent Statement

Before electrode implantation, all patients gave written informed consent for participation in research studies and for publication of the results. The patient SEEG data and clinical information were handled anonymously.

## 6 Data Availability Statement

Compliance with Italian governing laws and Ethical Committee regulations prohibits the sharing of raw data and patient details. However, interim data and final results that support the findings of this study can be obtained from the corresponding authors upon reasonable request.

## A Supplementary materials

### A.1 Supplementary methods

#### A.1.1 The bistability in Thom’s cusp catastrophe

Catastrophes emerging from complex systems are characterized by sudden and violent onsets, detrimental consequences, and are suggested to be generated by only few candidate mechanisms [43]. We suggest the emergence of epileptic hypersynchrony in neuronal population to be explain by Thom’s cusp catastrophe:

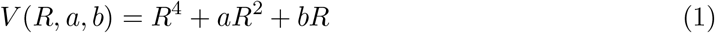

where *R* is synchrony; the stable solution for the potential function satisfies *V* :

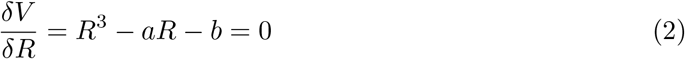

which defines the cusp fold, wherein the attractor of *R, e.g.*, experimentally observed mean synchrony over time, is controlled by two parameters: *a* modulates the degree of *R, e.g.*, a large *a* results in hypersynchrony; as *b* increases, bistability in *R* gradually emerges (Fig 1). High bistability invariably predicts catastrophic events across a wide array of complex systems (reviewed in [36]), and likely is the underlying bifurcation mechanism for epileptic seizures.

#### A.1.2 Bistability in a modified Kuramoto model

We built a modified Kuramoto model based on the classic definition [72] to investigate the catastrophic hypersynchrony in neuronal oscillation dynamics. The theory and motivation of the model were discussed in detail in [36]. Briefly, we introduced a state-dependent noise to this Kuramoto model to reflect the local-positive feedback. The model contained 200 all-to-all connected oscillators, and the dynamics of each oscillator *i* of the model is a scalar phase time series *θ*_*i*_(*θ* ∈ 0 : 2*π*) defined as:

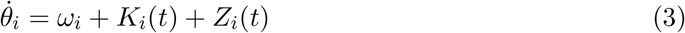

where, *ω*_*i*_ = 11 Hz is the natural frequency of the oscillators; *K*_*i*_ is the coupling between oscillators and the *Z*_*i*_ is the state-dependent noise. The coupling function *K*_*i*_ can be interpreted as oscillator *i* adjusts its phase due to the interaction with all other oscillators in the model and is defined as:

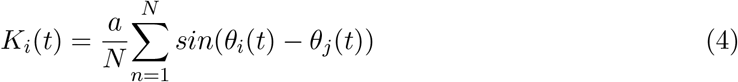

where *a* is the coupling strength between oscillators (Fig 1); *N* = 200 is the number of oscillators in the model. The noise *Z*_*i*_ is defined as:

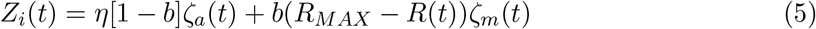

where *η* = 0.2 is a constant to weight the noise influence, *ζ*_*a*_(*t*) and *ζ*_*m*_(*t*) represent additive and multiplicative noise, respectively, and they are two independent Gaussian time series with zero mean and unit variance; the strength of positive local feedback *b* scales the influence of *ζ*_*m*_(*t*) depending on the current level of synchrony *R*; *R*_*MAX*_ = 0.96 is the maximal synchrony the population can reach, and the order parameter *R*, aka the synchrony among all the oscillators, is defined as:

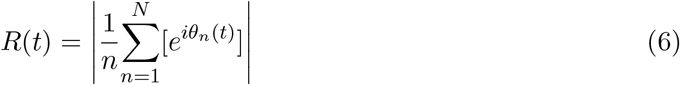

hence when the population is in full synchrony, *R* → 1; when the population is fully desynchronized, *R* → 0.

#### A.1.3 Criticality assessments

##### Assessing neuronal bistability with the BiS index

We used the BiS index to assess the bistability of a power time series (*R*^2^). Briefly, we followed the approach proposed in [39, 40] to obtain the probability distribution function (*pdf*) of a narrow-band power time series *R*^2^ with 200 bins. Next, because the square of a Gaussian process has an exponential *pdf*, we used maximum likelihood estimate (*MLE*) to fit the observed *pdf* with a single-exponential function:

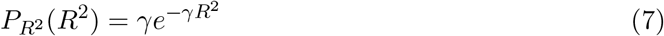

next, the same *pdf* was fitted with a bi-exponential function:

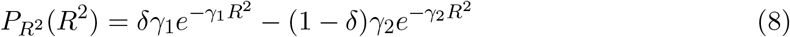

where where *γ*_1_, *γ*_2_ are the two exponents and *δ* is a weighting factor. Next, we used the Bayesian information criterion (BIC) to assess the fitting of eq.7 and eq.8:

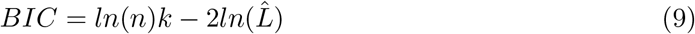

where, *n* is sample number; 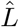 is the likelihood function; *k* is the number of parameters, i.e., for eq.7, *k* = 1 and for eq.8 and *k* = 3. A better fitted model yields a small *BIC* estimate. Hence, the *BIC* imposes a penalty to model complexity of eq.8 for two more degrees of freedom than eq.7. Finally, the BiS index is computed as the *log*_10_ transform of the difference between the *BIC* of the two models *dBIC* = *BIC*_*singleExp*_ −*BIC*_*biExp*_ as :

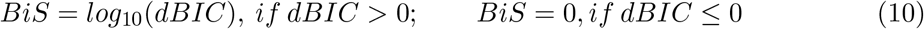

because a better fitted model yields a small *BIC* estimate, the *BiS* will be large if the biexponential model is a better model for the observed *pdf* of the oscillation power time series.

##### Assessing neuronal LRTCs using the DFA

We followed the approach proposed by [27] to assess the LRTCs of the narrow-band oscillation amplitudes with linear detrend fluctuation analysis (DFA). The theory, technical details, and the toolbox of the DFA are well described in [73]. Briefly, an estimated DFA exponent reflects the finite-size power-law scaling in narrowband amplitude fluctuations based on the assumption that the gradual evolution of a monofractal time series would result in a normal distribution where the fluctuations can be captured by the second order statistical moments. In practice, DFA characterizes how fast the overall root mean square of local detrend fluctuations *F* grows with increasing sampling window size *L*:

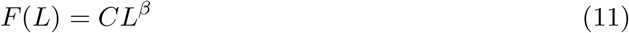

where *C* is a constant; *β* is the DFA exponent and *β >* 0.6 can be safely considered as a sign of a critical-like process, e.g., significantly greater than that of a random walk process (*DFA* = 0.5, see Fig 2 and Supplementary Fig 4 for fitting examples). Here, for assessing narrow-band amplitude DFA, we set *L* as 40 windows with their size ranging from 20-cycle length (*e.g.*, 2 seconds for 10 Hz) to 2.5 min with equal *log* distance between windows; the *F* (*L*_*i*_) were computed for a given window size *L*_*i*_ as the mean of the root mean square of the detrend fluctuation with 50% overlap between neighboring windows, and a window was excluded from the mean if more than 10% of its samples contain interictal epileptiform events; the linear regression of eq.11 was done with a bi-square fitting weighted by the square root of observed window number for a given *L*_*i*_.

##### Assessing neuronal excitation-inhibition with the fE/I

We followed the approach proposed by [29] to assess the functional excitation/inhibition (*E/I*) of a narrow-band oscillation amplitudes with the *f E/I* index:

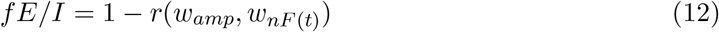

where *w*_*amp*_ denotes observed windowed oscillation amplitudes; *w*_*nF*_ (*t*) is the observed windowed detrend function of the amplitude-normalized signal profiles; *r*(*w*_*amp*_, *w*_*nF*_ (*t*)) denotes the Pearson’s correlation between *w*_*amp*_, *w*_*nF*_ (*t*). Thus, based on Bruining’s hypothesis that, if a time series is generated by a critical-like mechanism, we expect to observe, for an inhibitiondominated ensemble *f E/I* < 1; for an excitation-dominated ensemble *f E/I* > 1, and for an *E/I* balanced ensemble *f E/I* = 1 (illustrated in Supplementary Fig 2 j). Here, for assessing narrow-band amplitude *f E/I*, we set window size as 50-cycle length (*e.g.*, 5 seconds for 10 Hz) with 60% overlap between windows.

#### A.1.4 Phase synchrony assessment

The phase coupling between two narrow-band time series *A* and *B* (complex-valued) can be quantified as the phase-locking value as [74]:

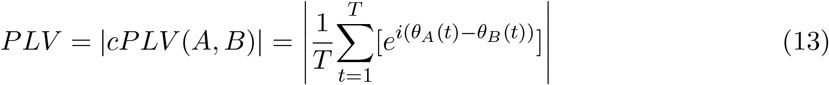

where *T* denotes the total number of independent samples; *θ*_*A*_ and *θ*_*B*_ are the instantaneous phases of *A* and *B*. We denote a *PLV* connectivity matrix as a “graph” *G*_*iP LV*_ = (*V, E*), where any SEEG contact *i* is a “node” *v*_*i*_ ∈*V* ; the *PLV* between two nodes (*v*_*i*_, *v*_*j*_) are edges, *e*_*k*_ = (*v*_*i*_, *v*_*j*_) ∈*E|v*_*i*_, *v*_*i*_ ∈*V .* The first- and second-order synchrony derivatives were computed using these *PLV* matrices.

### A.2 Supplementary results

#### A.2.1 Synchrony derivatives were robust against various random attacks

The anatomical coverage and number of SEEG contacts were variable across subjects. Moreover, some *PLV* edges were excluded from analyses if two contacts shared the same white-matter reference [46]. This raised a slight concern about whether the spatial sampling variability and missing edges in the *PLV* matrices would bias the first- and second-order Synchrony derivatives (Synch.D). We conducted random deletions to the edges and to the nodes in graphs and then compared Synch.D before and after the deletions to ask whether synchrony derivatives were resilient against these varying factors. This was done by computing the Pearson’s correlation *r* between the synchrony derivatives between the original and assaulted graphs, and as *r* → 1 means that the nodal topological features were preserved after the assault. Additionally, these Synch.D estimates were computed with weighted *PLV* graphs would be different after removing edges or nodes; in the SZ-classification analyses, we normalized synchrony derivatives within subjects, and therefore different estimates was not a concern here. After the shared-reference exclusion, subjects’ PLV graph density was 0.92 *±* 0.04(*mean ± std, range* : 0.81 − 0.96), where one means a fully connected network without nodal self-connection. To ascertain that the missing edges did not significantly impact the synchrony derivatives, we performed population-level edge deletion tests. We took 4.8 Hz *PLV* matrices from the 55 patients of the SZ-classification cohort. We next assaulted each subject’s PLV matrix by randomly deleting *x*% of edges in one surrogate run, and then compute the Pearson’s *r* between the Synch.D between the nodes from the original and assaulted graphs. We conducted 500 of such surrogates within each patient and across edge deletion ratio *x*% ∈ [10 : 10 : 90, 91 : 1 : 98], which assured that all synchrony derivatives (*i.e.*, We, EVC, Cc, and LE) were resilient to random edge deletion – specifically for the range of the missing data observed (Supplementary Fig 4 **a, c**).

Next, to ascertain that heterogeneity in SEEG spatial sampling or sub-sampling the targeted network did not significantly impact the synchrony derivatives, we performed population-level nodal removal tests. We used 4.8 Hz PLV matrices from the 55 patients and assaulted each subject’s *PLV* matrix by randomly removing *y*% of nodes in one surrogate run, and then compute the Pearson’s *r* between the synchrony derivatives of the remaining nodes from the assaulted and the original graphs. We conducted 500 of such surrogates within each patient and across nodal removal ratio *y*% ∈ [10 : 10 : 90, 91 : 1 : 95], which assured that all synchrony derivatives (*i.e.*, We, EVC, Cc, and LE) were resilient to random nodal removal (Supplementary Fig 4 **b, d**).

#### A.2.2 Supervised SZ-Classification

The individual variability in SZ-classification outcomes can be explained by several factors. Individual area under (AUC) the receiver operator characteristic curve was negatively correlated with the standard deviation of edge-distance (Spearman’s rank *r* = −0.3, *p <* 0.026), but was not correlated with mean edge distance (Spearman’s rank *r* = −0.24, *p >* 0.076) nor total number of edges (Spearman’s *r* = −0.23, *p >* 0.091), which means relatively concentrated spatial sampling rather than the spatial extent of the spatial sampling helps with increasing individual AUC. The *L*_*SZ*_ (0 − 100%, *i.e.*, the likelihood of being an SZ contact) assigned by the Random Forest classifier were different between the SZ and nSZ contacts (unpaired t-test, *p <* 6.8 *×* 10^−157^), and the difference in SZ and nSZ contacts’ EZ-likelihood predicted patient AUC (Spearman’s rank *r* = 0.93, *p <* 10^−6^, Supplementary Fig 7 k).

### A.3 Supplementary Figures

#### A.3.1 Supplementary Fig 1

**Figure 1.**
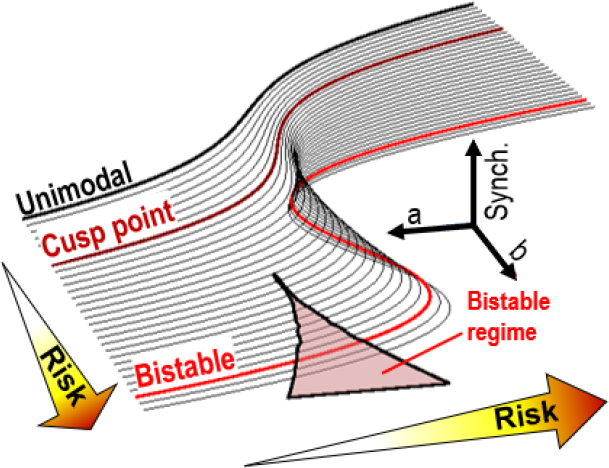
The cusp catastrophe offers a canonical explanation to the link between high bistability and catastrophic neuronal hypersynchrony. On the cusp fold, each point along the curve indicates the stable solution of *R* with a given pair of *a* and *b*; a large R corresponding to seizure-like hypersynchrony, and as *b* increases, the curves shifts shape from smooth to discontinuous transition in the temporal dynamics of *R*.

#### A.3.2 Supplementary Fig 2

**Figure 2.**
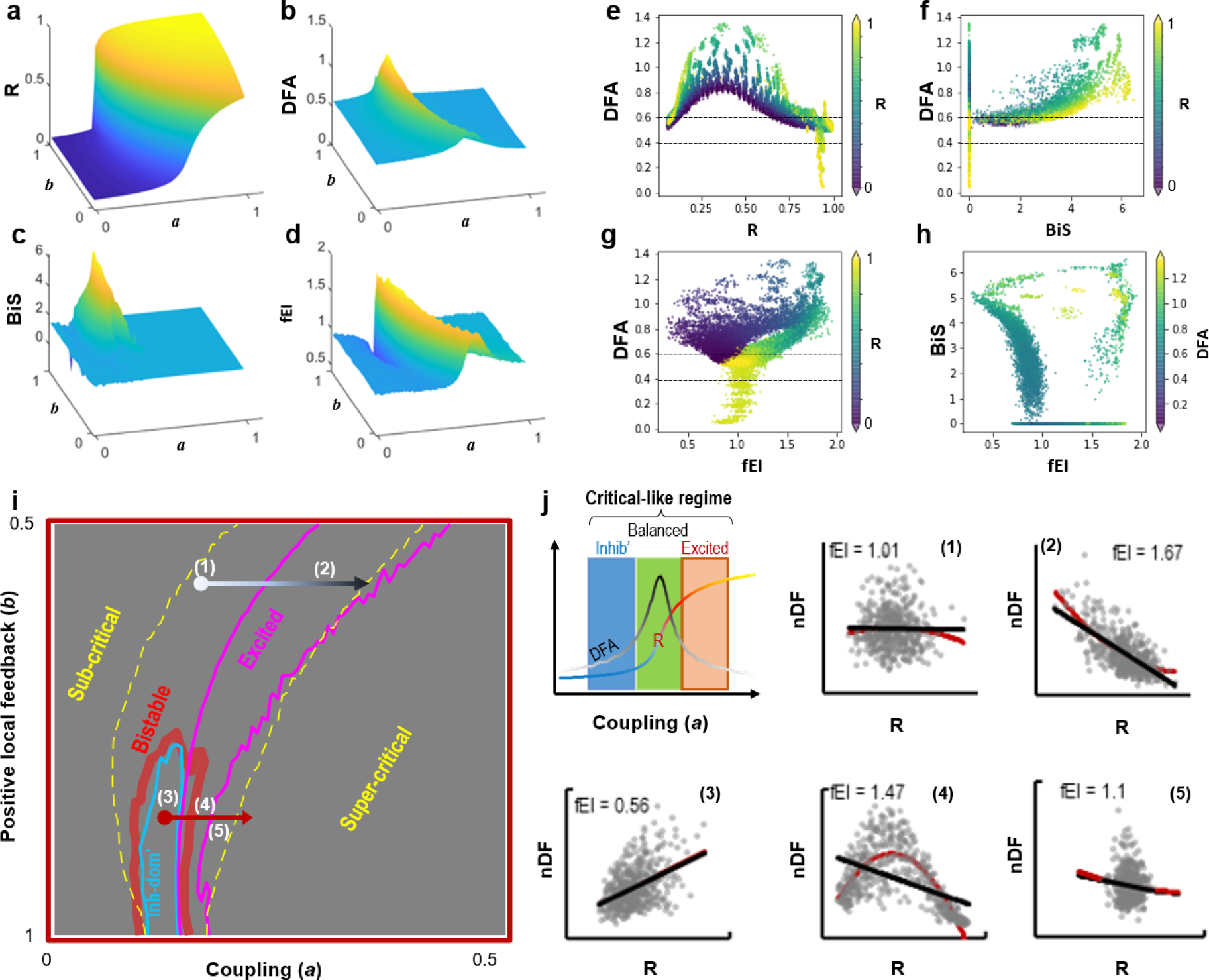
High bistability in neuronal synchrony of the model indicates a shift toward catastrophic hypersynchrony. **a–d.** Assessments averaged over 20 independent model realization for **(a)** model synchrony (*R*), **(b)** DFA exponent **(c)** BiS index and **(d)** fE/I index. **e–h.** Co-variability between the criticality assessments. **i.** In high bistability regime, a small increment in *a* (red vector) could drive the model from adaptive critical-like dynamics into seizure-like hypersynchrony, during which the model can show radical switching from inhibition-dominance (3) to excitation-dominance (4). **j.** Schematic showing how the fE/I index utilizes distinct relationship between *R* and *nDF* (normalized detrend function) to capture the model operation dynamics in three regions inside critical regime. Inhib’: inhibition-dominance; excited: excitation-dominance; (1–5) the functional excitation-inhibition as response to *a* increment inside and outside of the bistable regime (as marked in (**i**)), each marker indicates the estimates of an analysis window of 1000 samples; also note that data can sometimes be fitted with a quadratic function indicating a wider operation region than one of the three regions.

#### A.3.3 Supplementary Fig 3

**Figure 3.**
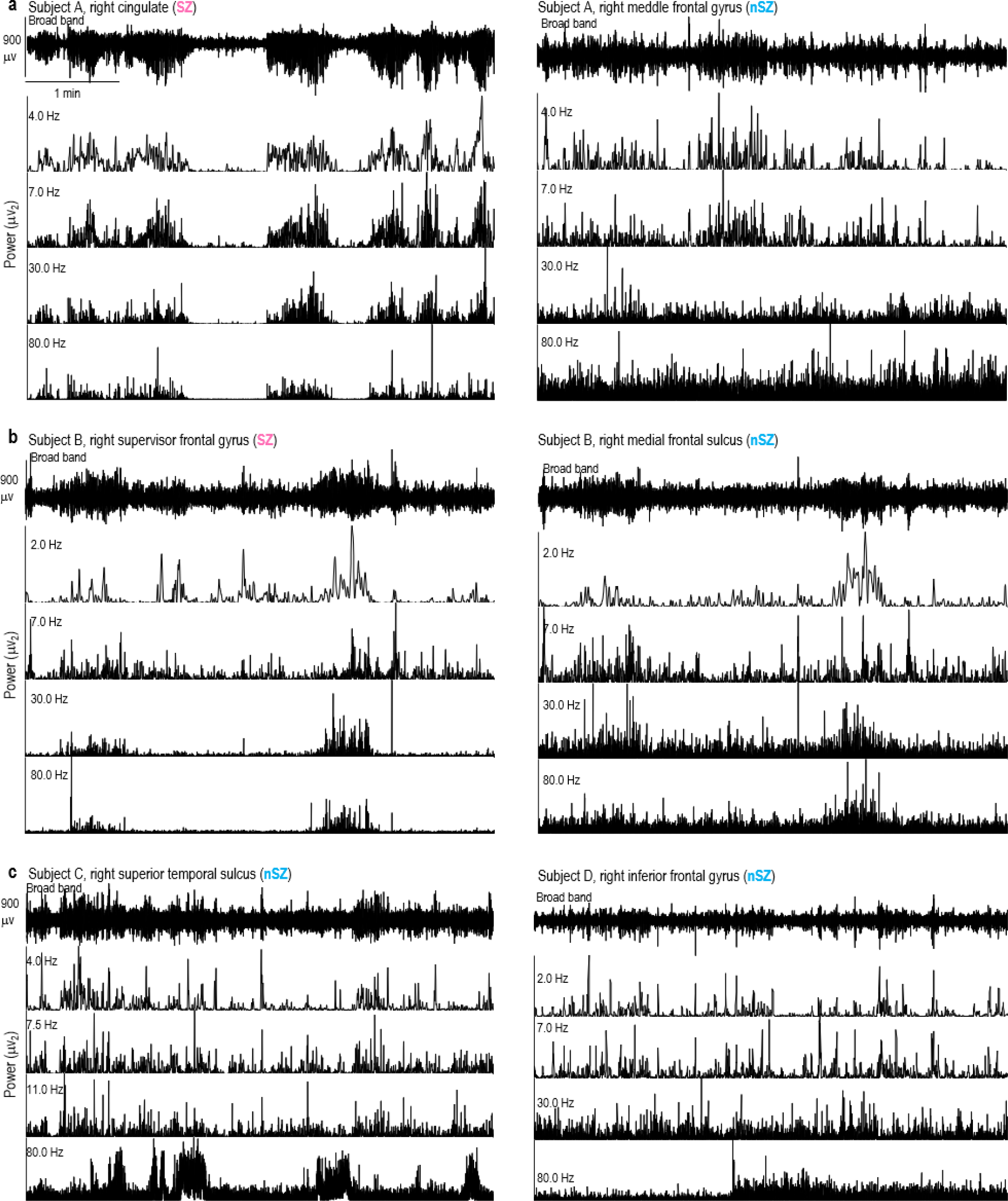
SEEG contacts from both SZ and nSZ can show bistable activity. **a–b.** Broad-band traces and narrow-band power time series showing that higher bistability in an SZ than in a nearby nSZ contact in two subjects. **c.** nSZ contacts can also demonstrate bistability as shown here in 4 and 80 Hz narrow-band oscillations from another two subjects.

#### A.3.4 Supplementary Fig 4

**Figure 4.**
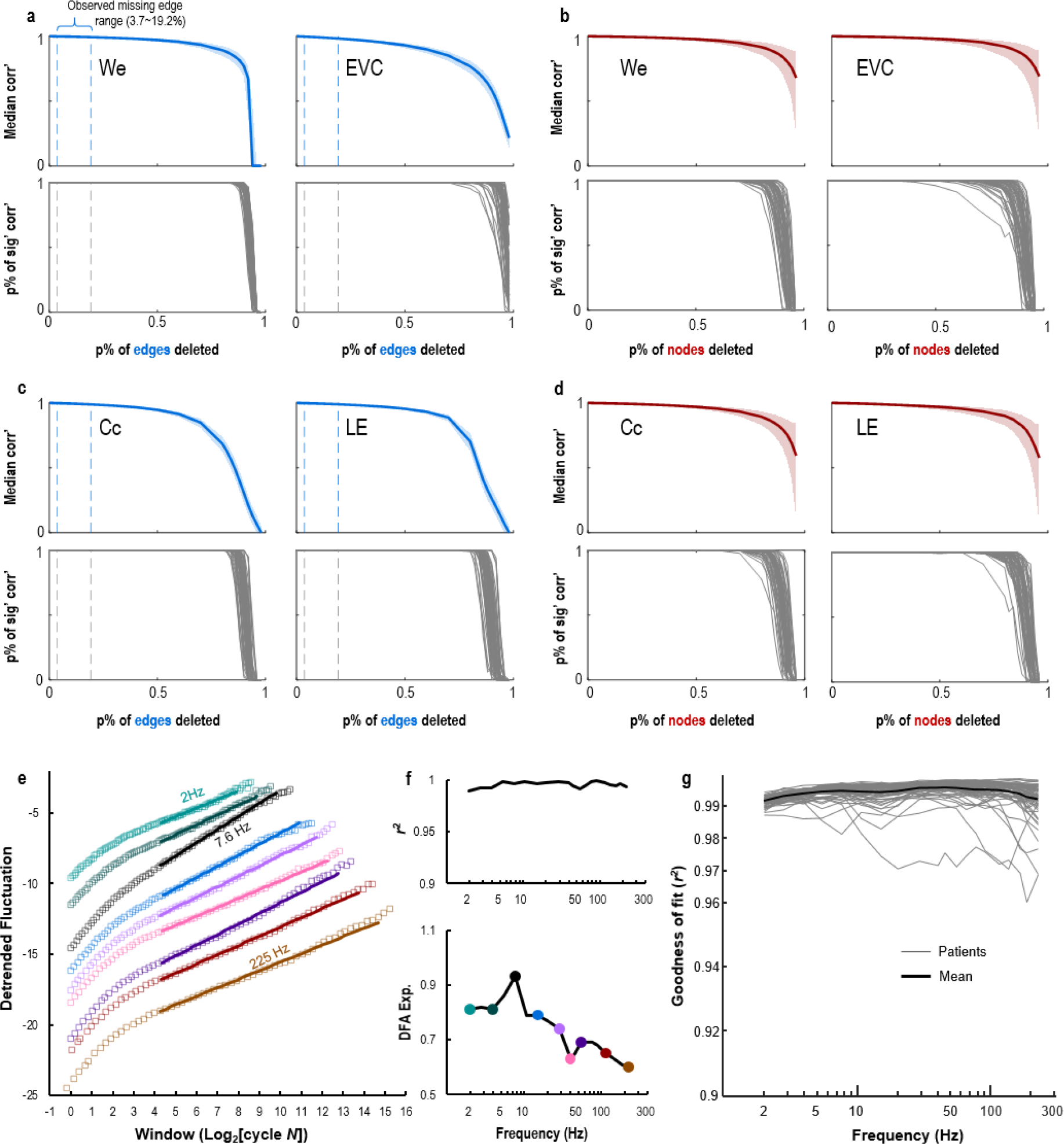
Synchrony derivatives were resilient against missing data, *e.g.*, under-sampling, and DFA goodness-of-fit was consistent across frequencies and subjects . **a–b.** The first- and **(c–d)** second-order synchrony derivatives (Synch.D) employed in SZ-classification were highly resilient to random edge deletion (blue) or nodal removal (red). ***Top:***median correlation (Pearson’s *r*) between the Synch.D of the original PLV graphs and the graphs under increasing levels of random assaults, computed over 500 iterations per subject for 55 subjects of the EZ-classification cohort; shaded areas indicate interquartile distance. ***Bottom***: corresponding percent of significant correlations (*p <* 0.01, FDR corrected) out of the 500 surrogates in each subject (thin lines). **e.** An example of how DFA fitting was done across narrow-band frequencies from a SEEG contact of a randomly selected subject. Markers indicate observed data. Lines indicate DFA linear fitting, and the fitting range was from 20 cycle-length of a given narrow-band frequency up to 25% of the 10-min resting-state recording. **f. *Top***: goodness of fit (*r*^2^) and ***bottom***: the DFA exponents of the contact from (e). **g.** DFA goodness-of-fit of individual patients’ mean across contacts (thin) and group mean (thick).

#### A.3.5 Supplementary Fig 5

**Figure 5.**
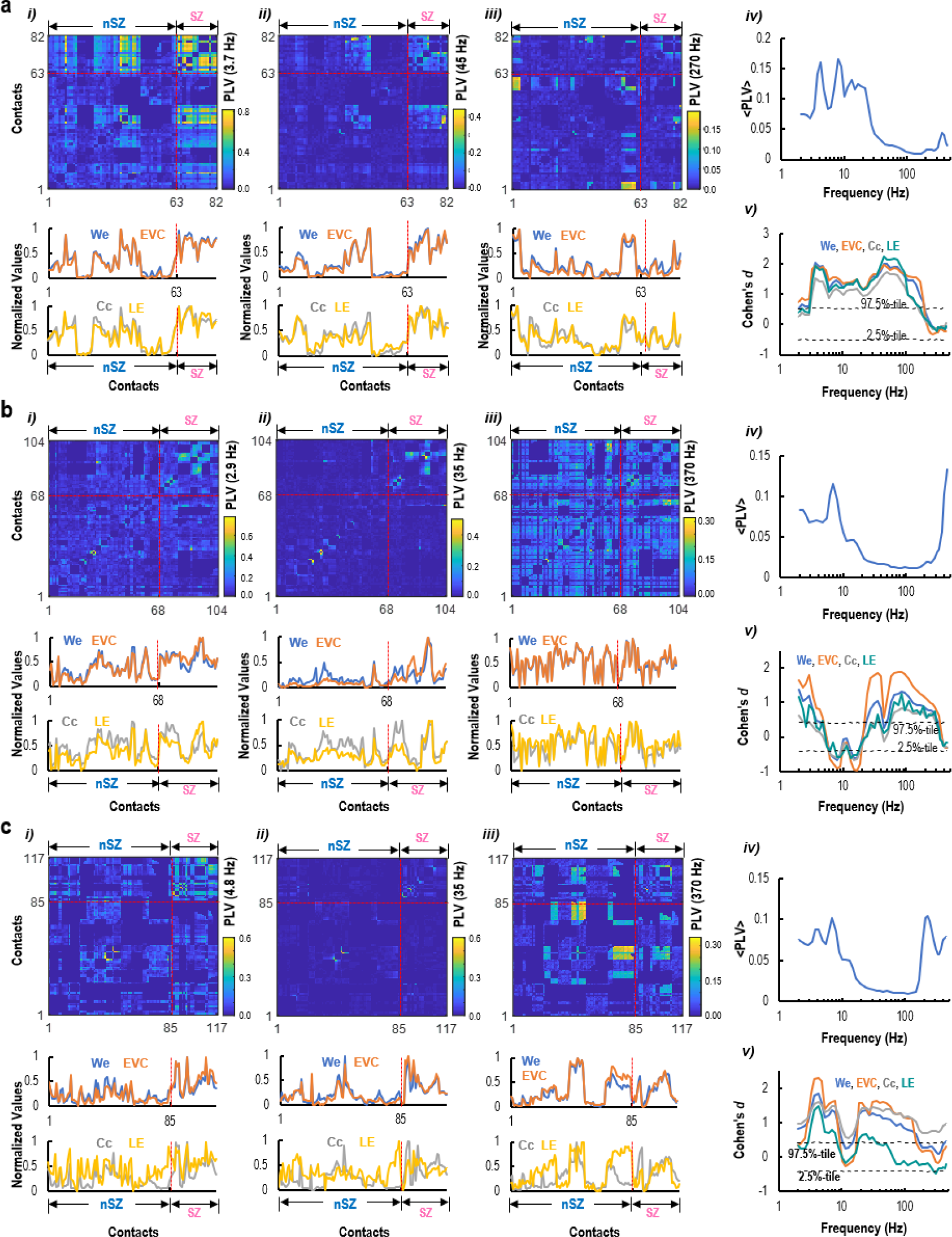
Individual level evidence of differences between SZ and nSZ in synchrony derivatives. **a–c.** The narrow-band frequency *PLV* matrices and their corresponding synchrony derivatives (Synch.D), and individual level differences between SZ and nSZ in three subjects. Within subjects: ***i-iii***) ***Top***: narrow-band *PLV* matrices; ***bottom***: *We* and *EV C* (first-order) and *Cc* and *LE* (second-order Synch.D) of the corresponding *PLV* matrices. ***iv***) Individual connectivity spectrum, *i.e.*, effective mean *PLV* edges across narrow-band frequencies. ***v***) Differences between SZ and nSZ in Synch.D.

#### A.3.6 Supplementary Fig 6

**Figure 6.**
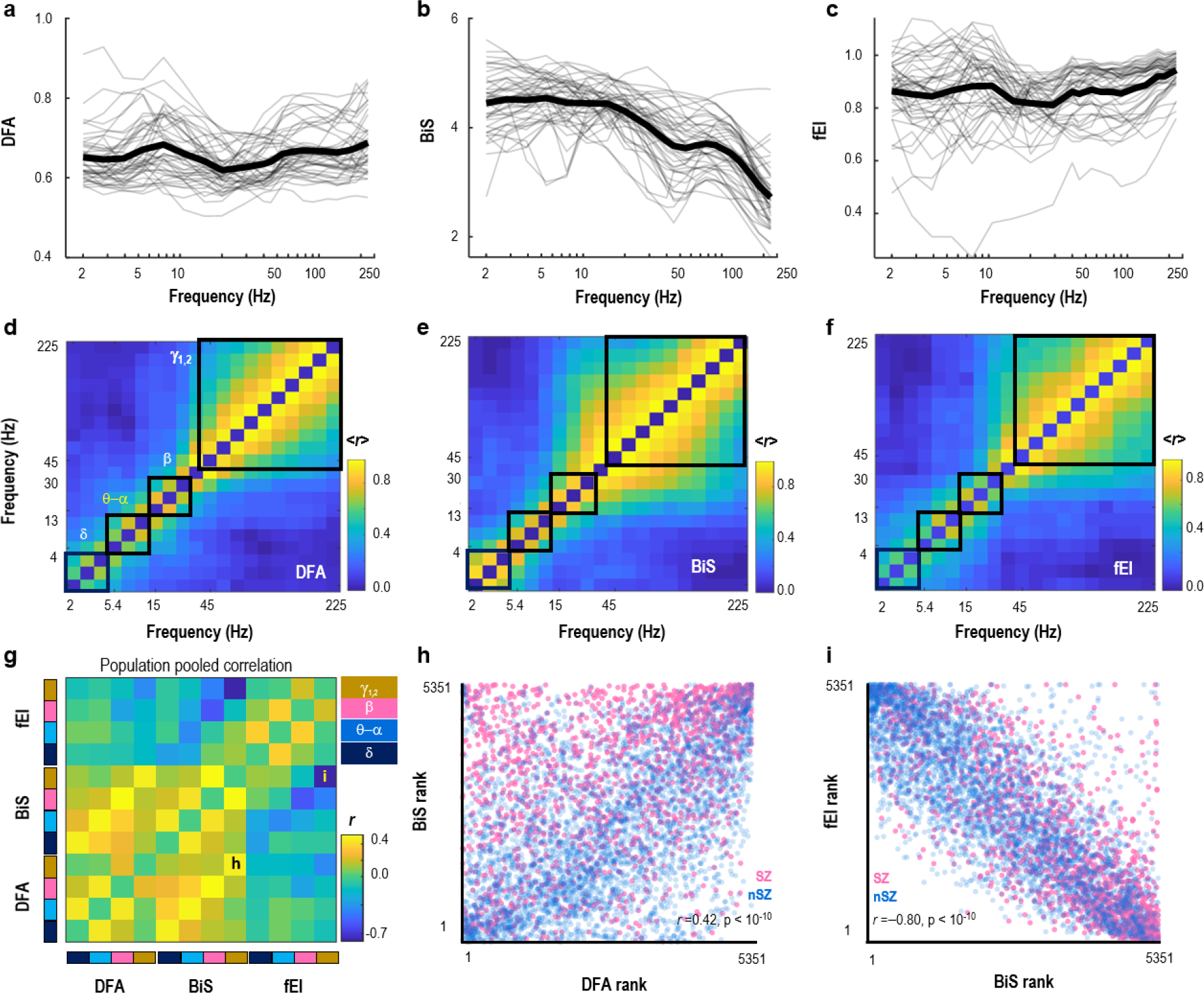
Supplementary information for frequency clustering by topological similarity. **a–c** Within patient mean (thin) and cohort average (thick) (**a**) DFA, (**b**) BiS, and (**c**) fE/I. **d–f** Cross-frequency topological similarity (Spearman’s rank *r*) for (**d**) DFA, (**e**) BiS, and (**f**) fE/I. **g.** Population pooled Spearman’s rank *r* between frequency clustered criticality assessments (validation for Fig 4 a), **h–i.** Example of the relationship between the Pearson’s rank of contact **(h)** *γ*_1,2_ band DFA and BiS estimates, and **(i)** *γ*_1,2_ band BiS and fE/I estimates, as indicated in **(g)**.

#### A.3.7 Supplementary Fig 7

**Figure 7.**
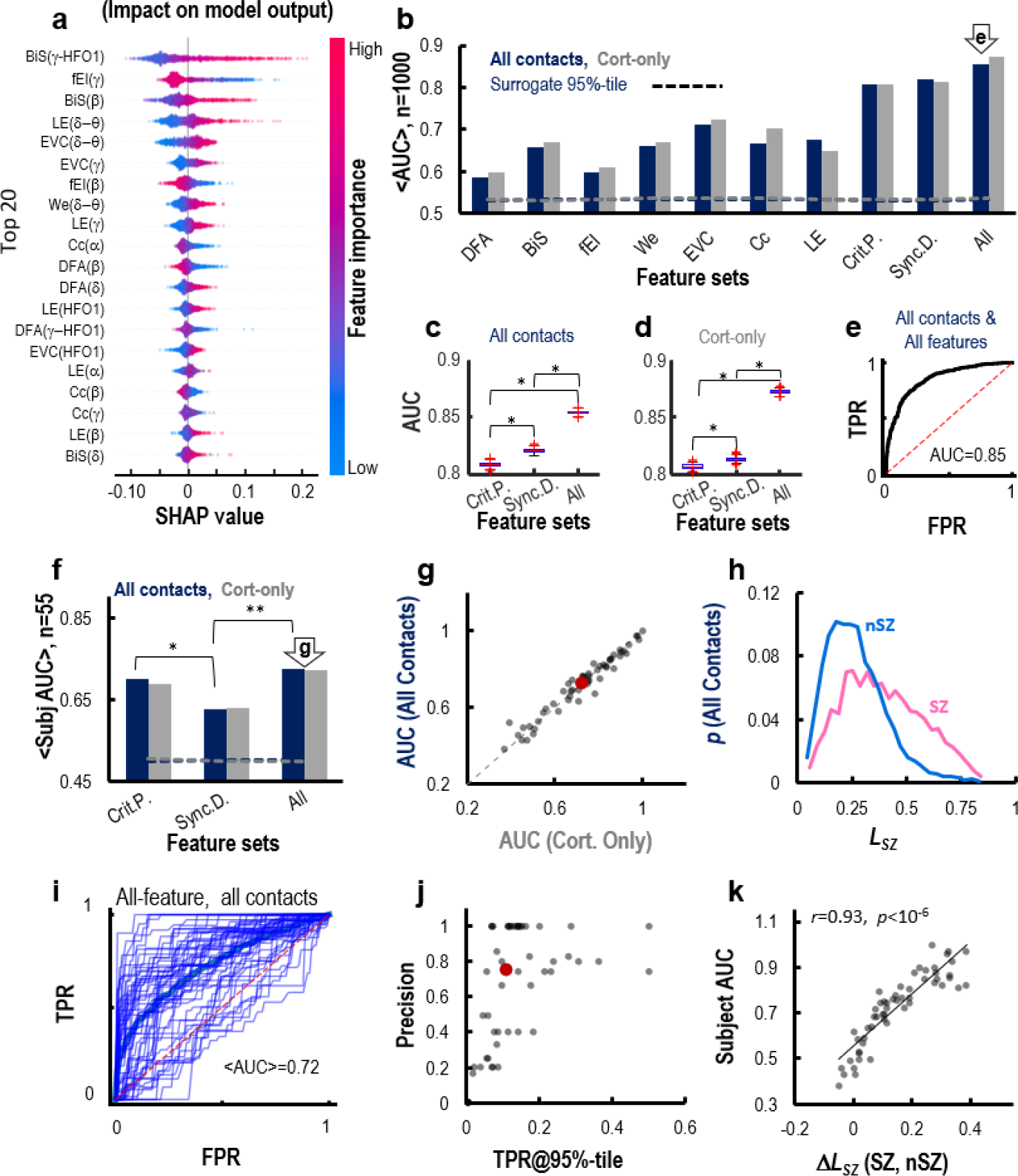
Supplementary information for the supervised SZ-classification using Random Forest (RF) algorithm. **a.** Importance ranking of the individual features assessed using the SHAP value. **b–e.** The support information for cohort level cross-validate. **b.** SZ-classification accuracy assessed using the area under (AUC) the receiver operating characteristic (ROC) for the RF classifier trained on different feature sets using all contacts (blue) and with subcortical contacts excluded (gray). **c–d.** The AUC differences between different feature combinations. **e.** The ROC of a randomly selected SZ-classification iteration when using “all features” and “all contacts”. **f–k.** Support information for within-patient SZ-classification. **f.** Difference between individual AUC (*, ** indicate *p <* 0.05 and 0.01). **g.** No difference between the individual AUC of SZ-localization trained on cortical-only and all contacts. **h.** Pooled probability distribution of the EZ-likelihood *L*_*SZ*_assessed by the RF for SZ and nSZ contacts. **i.** Individual (thin) and group mean (thick) ROC curves. **j.** Individual (gray) and mean (red) precision as a function of true positive rate (TRP) when *L*_*SZ*_threshold was held at 95% confidence interval for SZ-classification using all features and all contacts. **k.** The individual AUC of the SZ-localization as a function of the within-subject difference in *L*_*SZ*_between SZ and nSZ.

#### A.3.8 Supplementary Fig 8

**Figure 8.**
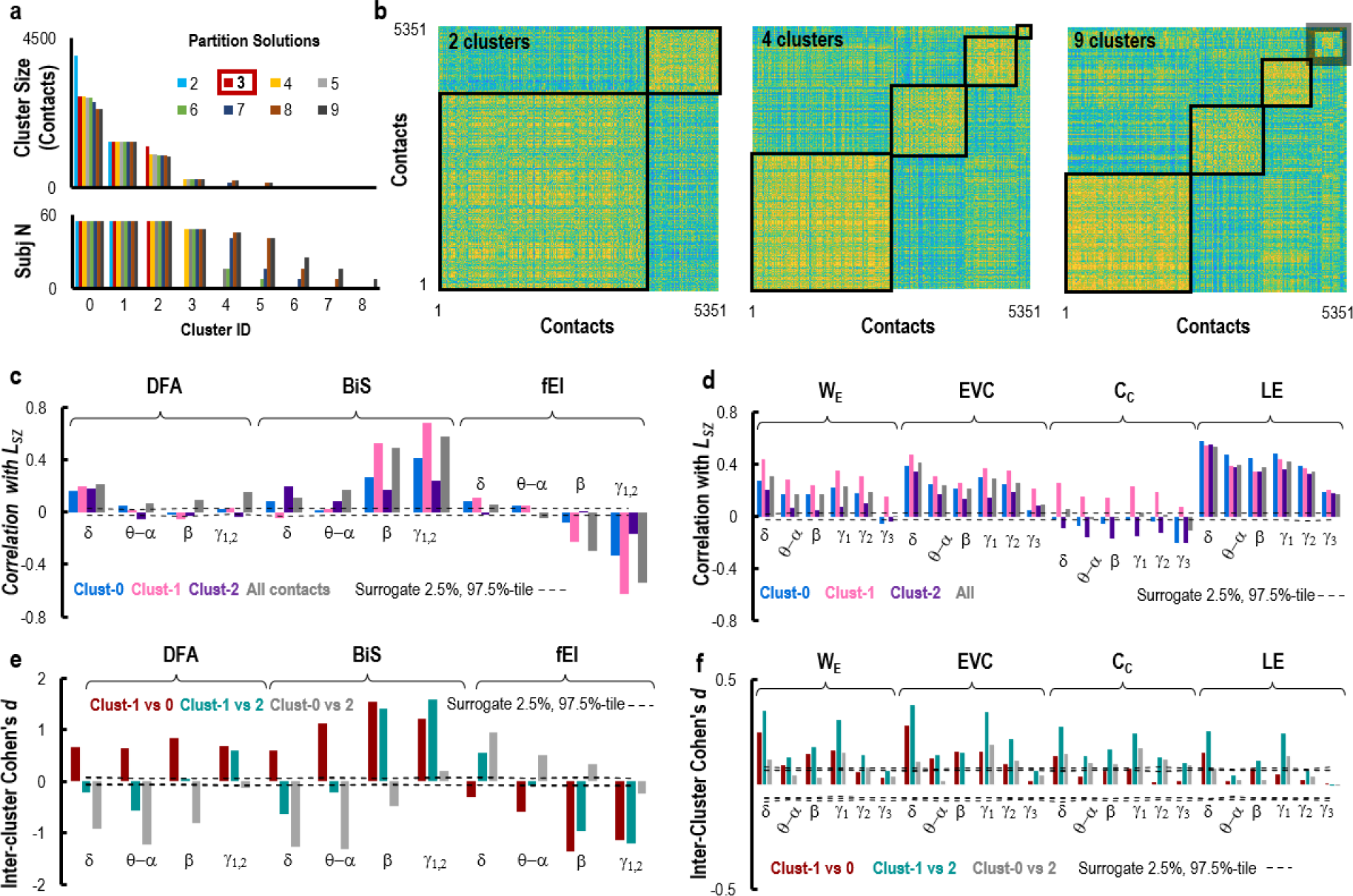
Supplementary information for unsupervised classification. **a.** Top: cluster size distribution; bottom: distinct subject number within each cluster across partition solution from 2 to 9 clusters color-coded. The 3-cluster solution was used in (Fig 4 e–i). **b.** Visualized partition solutions of the inter-contact similarity (Spearman’s rank *r*) matrix for 2-, 4-, and 9-cluster solutions. The gray box in the 9-cluster solution contains 6 small clusters. **c–d.** For the 3-cluster solution, the correlations between the *L*_*SZ*_and **(c)** criticality assessments and **(d)** synchrony derivatives. **e–f.** The inter-cluster contact difference in **(e)** criticality assessments and **(f)** Synchrony derivatives. Dashed lines in **(c-f)** indicate confidence intervals (*α <* 0.05) observed from 10^4^ surrogates.

