## Supplementary material for "Neuronal Synchrony and Critical Bistability: Mechanistic Biomarkers for Localizing the Epileptogenic Network": Subject information.

| ID | Seizure Zone | Age | Sex | Medication | Outcome (Engel score) |
| --- | --- | --- | --- | --- | --- |
| 1 | Right mesial frontal | 21 | M | Carbamazepine 600mg, Levetiracetam 1500mg | IB (8 years) |
| 2 | Right temporal insular | 26 | M | Carbamazepine 1200mg, Primidone 750 mg, Clonazepam 10mg | IA (37 months) |
| 3 | Right temporal | 38 | M | Carbamazepine 1200mg, Dilantin 450mg, Lacosamide 300mg, Clonazepam 10mg | No surgery |
| 4 | Left temporo-parietal | 38 | F | Phenobarbital 100mg, Topiramate 100mg, Levetiracetam 3000mg | IA (25 months) |
| 5 | Left temporal-insular | 24 | M | Oxcarbazepine 600mg, Lacosamide 400mg | IA (15 months) |
| 6 | Right temporo-insular | 40 | F | Levetiracetam 1000mg, Lacosamide 350mg, Sertraline 50mg, Lorazepam 1mg | IIIA (32 months) |
| 7 | Precuneus | 19 | M | Oxcarbazepine 600mg, Lacosamide 400mg | No surgery |
| 8 | Functional epilepsy | 20 | F | Carbamazepine 1200mg, Levetiracetam 1500mg | No surgery |
| 9 | Right occipito-temporo-parietal | 20 | M | Lamotrigine 400mg, Levetiracetam 3000mg, Lacosamide 400mg | IA (3 years) |
| 10 | Left temporal | 35 | M | Topiramate 200mg, Carbamazepine 900mg | IA (12 months) |
| 11 | Right temporal anterior | 28 | M | Carbamazepine 1200mg | IIA (66 months) |
| 12 | Temporo-hip | 36 | F | Carbamazepine 1400mg, Levetiracetam 3000mg | IIA (6 months) |
| 13 | Left frontal anterior | 40 | M | Levetiracetam 2750mg, Carbamazepine 800mg, Primidone 750mg | IA (12 months) |
| 14 | Thermo-coagulation multiple sites | 39 | M | Oxcarbazepine 1800mg, Clobazam 20mg | IA (36 months) |
| 15 | Right fronto-temporo-insular | 24 | F | Carbamazepine 1000mg, Clobazam 20mg, Lomotrigine 200mg | IVA (24 months) |
| 16 | Left parieto-opercolo-insular | 31 | F | Carbamazepine 1200mg, Clobazam 40mg, Phenobarbital 75mg | IVA (12 months) |
| 17 | Right temporo-perisilvian | 34 | F | Phenobarbital 150mg, Lacosamide 400mg, Clobazam 10mg | IIC (38 months) |
| 18 | Right perisilvian-insular | 17 | M | Carbamazepine 800 mg, Lamotrigine 400mg | IVA (26 months) |
| 19 | Right temporo-parieto-occipital | 36 | M | Oxcarbazepine 1200mg, Phenobarbital 150mg, Valproate 1000mg | IA (35 months) |
| 20 | -- | 32 | F | Carbamazepine 700mg | No surgery |
| 21 | Left temporal antero-mesial | 32 | M | Carbamazepine 1200mg, Levetiracetam 750mg | IA (61 months) |
| 22 | Right fronto-centro-insular | 33 | M | Carbamazepine 800mg, Lacosamide 800mg, Zonisamide 250mg | IIA (38 months) |
| 23 | Left temporal | 21 | F | Levetiracetam 1750mg, Lacosamide 400mg, Valproate1000mg | IA (24 months) |
| 24 | Right parietal | 23 | M | Levetiracetam 3000mg, CBZ 1000mg, Lacosamide 500mg | IA (24 months) |
| 25 | Thermo-coagulation | 46 | M | Carbamazepine 1200mg, Phenobarbital 100mg | IA (24 months) |
| 26 | Right frontal | 20 | F | Valproate 800mg, Clobazam 10mg | IIA (36 months) |
| 27 | Right fronto-mesial | 21 | M | Carbamazepine 800mg, Levetiracetam 3000mg, Nitrazepam 1.5mg | IIIA (13 months) |
| 28 | Right fronto-central | 22 | M | Lamotrigine 400mg, Levetiracetam 2000mg | IA (24 months) |
| 29 | Right frontal | 20 | M | Carbamazepine 600mg, Rufinamide 1500mg | IVA (13 months) |
| 30 | Right frontal | 44 | F | Carbamazepine 1200mg, Zonisamide 400mg, Phenobarbital 1000mg | IC (24 months) |
| 31 | -- | 17 | M | Carbamazepine 300mg | No surgery |
| 32 | -- | 14 | M | Levetiracetam 1500 mg, Clobazam 5mg | No surgery |
| 33 | Right temporal antero-mesial | 30 | F | Oxcarbazepine 2000mg, Phenobarbital 150mg | IIA (36 months) |
| 34 | -- | 24 | M | Carbamazepine 16000mg, Levetiracetam 4000mg | No surgery |
| 35 | -- | 29 | F | Levetiracetam 3000mg | No surgery |
| 36 | Right orbito-temporal | 29 | F | Zonisamide 400mg, Levetiracetam 750mg, Carbamazepine 1400mg | IA (62 months) |
| 37 | -- | 45 | F | Lacosamide 500mg, Valproate 1000mg, Zonisamide 200mg | No surgery |
| 38 | Thermo-coagulation multiple sites | 34 | F | Carbamazepine 1000mg, Levetiracetam 2500mg | IA (12 months) |
| 39 | Thermo-coagulation multiple sites | 50 | M | Levetiracetam 2000mg, Lacosamide 600mg | IIA (6 months) |
| 40 | Left occipital | 17 | F | Carbamazepine 1200mg, Levetiracetam 1500mg, Lacosamide 300mg | IB (49 months) |
| 41 | Right temporal | 44 | F | Topiramate 300mg, Oxcarbamazepine 1200mg | IIA (50 months) |
| 42 | -- | 27 | M | Carbamazepine 800mg, Lamotrigine 200mg | No surgery |
| 43 | -- | 46 | M | Carbamazepine 1200mg, Levetiracetam 3000mg, Lacosamide 150mg, Clobazam 20mg | No surgery |
| 44 | Right antero-frontal | 28 | M | Carbamazepine 1000mg, Levetiracetam 1000mg | IIIA (61 months) |
| 45 | Thermo-coagulation right temporo-parieto-perisilvian | 27 | F | Topiramate 200mg, Lamotrigine 200mg | IA (5 years) |
| 46 | Right temporal antero-mesial | 42 | F | Lacosamide 500mg | IB (36 months) |
| 47 | Left parieto-temporal | 15 | M | Carbamazepine 900mg | IA (6 months) |
| 48 | Thermo-coagulation right temporo-opercular | 37 | M | Carbamazepine 900mg, Levetiracetam 3000mg | IVA (12 months) |
| 49 | Left frontal | 30 | F | Carbamazepine 1200mg, Lamotrigine 200mg, Clobazam 20mg | IA (5 years) |
| 50 | Left frontal | 15 | F | Levetiracetam 1250mg, Oxcarbamazepine 1200mg | IA (4 years) |
| 51 | Thermo-coagulation | 41 | M | Levetiracetam 3000mg, Lacosamide 400mg | IA (2 years) |
| 52 | Right temporo-occipital | 37 | M | Lamotrigine 600mg, Levetiracetam 2000mg | IA (2 years) |
| 53 | Right temporal | 29 | M | Carbamazepine 1400mg, Levetiracetam 3000mg, Clobazam 10mg | IA (31 months) |
| 54 | Left opercolo-insular | 10 | F | Carbamazepine 800mg | IIIA (34 months) |
| 55 | Right temporo-frontal | 40 | F | Lamotrigine 600mg, Clobazam 20mg, Phenytoin 500mg | IA (13 months) |
| 56 | Left temporo-insular-operculum | 29 | F | Carbamazepine 1800mg, Clobazam 20mg | IA (24 months) |
| 57 | Right temporal | 27 | M | Lamotrigine 400mg, Topiramate 400mg | IA (24 months) |
| 58 | -- | 26 | M | Phenytoin 400mg, Topiramate 500mg | No surgery |
| 59 | Left temporal | 17 | M | Oxcarbamazepine 1500mg, Clobazam 20mg, Levetiracetam 2500mg | IA (36 months) |
| 60 | Right temporo-mesial | 25 | F | Topiramate 75mg, Carbamazepine 1500mg | IA (12 months) |
| 61 | Nodular heterotopia | 24 | F | Carbamazepine 1000mg, Levetiracetam 500mg, Clobazam 20mg | IVA (12 months) |
| 62 | Left temporo-perisilvian | 37 | F | Carbamazepine 1200mg, Lamotrigine 550mg | IIIA (12 months) |
| 63 | Left temporal antero-mesial | 32 | F | Clobazam 20mg, Phenobarbital 45mg | IIA (55 months) |
| 64 | Right temporo-occipital | 44 | M | Carbamazepine 800mg, Levetiracetam 3000mg, Phenobarbital 125mg | IA (24 months) |

**Supplementary Table 1 *Demographic data for the SEEG patient cohort***. Thermo-coagulation refers to the practice used in surgical intervention of drug-resistant focal epilepsy where current is injected in a bipolar derivation in order to increase the peri-contact temperature. Dashed entries in EZ locations mean that no single focal location was identified. Age column represent the age of the patients at the recording date. Drugs are reported with their active principle names. Drug dosage is expressed milligrams and refers to the morning dosage measured at the day of the recording. Outcome is expressed as Engel scores and number in parenthesis refer to the point time after surgery when the visit occurred.
